# IL-17A/IFN-γ producing γδ T cell functional dichotomy impacts cutaneous leishmaniasis in mice

**DOI:** 10.1101/2024.01.22.576494

**Authors:** Júlio Souza dos-Santos, Luan Firmino-Cruz, Diogo Oliveira-Maciel, Alessandra Marcia da Fonseca-Martins, Tadeu Diniz Ramos, Letícia Nunes-Souza, Rodrigo Pedro Soares, Daniel Claudio Oliveira Gomes, José Mengel, Bruno Silva-Santos, Herbert Leonel de Matos Guedes

## Abstract

γδ T cells are innate-like lymphocytes with pleiotropic roles in immune responses to pathogens, often ascribed to their IL-17A-producing or IFN-γ-producing γδ T cell subsets. Here we investigated the impact of this functional dichotomy on cutaneous leishmaniasis, a set of neglected diseases caused by parasites of the *Leishmania* genus. We demonstrate that in Sv129 mice susceptible to *Leishmania amazonensis*, Vγ4^+^ γδ T cells are the main source of IL-17A. Furthermore, in type 1 interferon receptor-deficient (A129) mice presenting increased susceptibility to infection, there is a higher frequency of IL-17A-producing γδ T cells when compared to wild-type mice. Mechanistically, we demonstrate that lipophosphoglycan (LPG) of *L. amazonensis* induces IL-17A-producing γδ T cells. Importantly, C57Bl/6 mice deficient in γδ T cells or in IL-17 receptor (IL-17RA) show reduced lesion sizes, consistent with a pathogenic role of IL-17A-producing γδ T cells in cutaneous leishmaniasis. Conversely, the adoptive transfer of FACS-sorted γδ T cells led to an accumulation of IFN-γ-producing γδ T cells in various susceptible strains of mice which associated with control of lesion development. These data demonstrate a pathophysiological dichotomy in which IL-17A-producing γδ T cells promote pathogenesis, whereas IFN-γ-producing γδ T cells display therapeutic potential in cutaneous leishmaniasis.

## Introduction

Cutaneous leishmaniasis (CL) is a neglected disease caused by protozoa of the genus *Leishmania*. It affects millions of people in the world, mainly low-income people, spreading in tropical and subtropical areas, and being found in 98 countries in Europe, Africa, Asia and the Americas^1^. Currently, there are more than 18 different species of *Leishmania* described as pathogenic to humans ^1,2^. Although likely underreported, current estimates of CL incidence range from 700,000 to 1.2 million cases per year^3^. Even though CL is the most frequent manifestation of *Leishmania* infection, visceral leishmaniasis (VL) can be fatal if left untreated^4^.

Infection by *Leishmania* spp. has been discussed for a long time as a trademark of the Th1/Th2 paradigm since the response of CD4^+^ T helper 1 (Th1) lymphocytes leading to the production of IFN-γ in C57BL/6 mice, or the CD4^+^ Th2 response leading to the production of IL-4 in BALB/c mice, culminate in resistant or susceptible phenotypes, respectively, in *L. major* infection^5^. The signaling induced by IFN-γ and T-bet is crucial for promoting the Th1 profile and inhibiting the Th2 profile through the downregulation of GATA-3^6^. Moreover, T-bet^-/-^ mice are more susceptible to *L. major* infection, consistent with a crucial role for Th1 cells in host protection from this parasite species ^7^.

In infections by *L. amazonensis* the dichotomy of susceptible and resistant mice is not evident, as a mixed of Th1/Th2 response is generated against the parasite, resulting in the production of both IFN-γ and IL-4 ^8,9^. In general, the production of IFN-γ induces control of the lesion and parasite load^10,11^, however, in *L. amazonensis* infection, CD4 T cells are overtly pathogenic^12^. Furthermore, *L. amazonensis* can subvert the immune response by inducing suppression of in macrophages^13–15^ and dendritic cells ^16^. *L. amazonensis* infection is also capable of inducing the expression of exhaustion markers such as PD-1 on CD4^+^ and CD8^+^ T cells, and PD-L1 on dendritic cells^17^. Treatment with anti-PD1 and anti-PD-L1 monoclonal antibodies restores the ability of T cells to produce IFN-γ, in addition to reducing the levels of IL-4 and TGF-β^17^.

The mixed Th1/ Th2 responses seen in *L. amazonensis*-infected mice are similar to those observed in human infections^18^. However, whereas in mice the Th2 response associates with disease^19^, in human leishmaniasis the production of cytokine transcripts of the Th2 profile was not necessary for disease maintenance in individuals infected by *L. amazonensis*^20^, suggesting that other cells may dictate susceptibility in human CL.

IL-17 derived from Th17 cells has been associated with the pathogenesis of *L.major* infection in BALB/c mice^21^, however, it has no impact on C57BL/6 mice^22^. IL-17^+^ cells have been associated with DCL caused by *L. amazonensis*^23^, but the underlying main source of this cytokine has not yet been revealed. Interestingly, in human leishmaniasis, the clinical presentations mediated by *L. amazonensis*, *L. major* and *L. braziliensis* infection associate with increased γδ T cells^24–30^.; and in experimental leishmaniasis, *L. amazonensis* infection in susceptible Sv129 mice induces the expansion of IL-17A producing γδ T cells, but not Th17 cells^31^.

IL-17 production by γδ T cells protects against fungi^32^ and bacteria, such as *Listeria* or *Staphylococcus*^33^. On the other hand, IFNγ-producing γδ T cells are particularly important in response to viral infections such as cytomegalovirus^34^. However, opposing effects of these cytokines, cytotoxic and wound healing functions can dictate the prognosis of various cancers^35,36^ and the outcome of parasitic diseases such as malaria^37^. In this context, IL-17 versus IFN-γ producing γδ T cells could also impact cutaneous leishmaniasis. Consistent with this hypothesis, the IFN-γ-mediated protection against *L. amazonensis* infection induced by oral LaAg vaccination was compromised upon depletion of γδ T cells^38^. However, a thorough investigation of the participation of distinct γδ T cell subsets in *L. amazonensis* infection is still lacking.

In this study, we characterized the γδ T cell population responding to *L. amazonensis* and, using transgenic mice, evaluated the functional role of γδ T cell subsets. We demonstrated that in Sv129 mice, which were susceptible to *L. amazonensis*, γδ T cells were the main source of IL-17A. These cells expanded more in type 1 IFNR-deficient mice, correlating with larger lesions and higher parasite loads. Additionally, in C57BL/6 mice, the deficiency of γδ T cells or IL-17 associated with smaller skin lesions. However, the transfer of γδ T cells to different mouse models resulted in the control of cutaneous leishmaniasis caused by *L. amazonensis* which associated with increased frequency of IFN-γ producing γδ T cells. Thus, we reveal a dichotomy of γδ T cells making IL-17A or IFN-γ that greatly impacts the outcome of *L. amazonensis* infection in mice. These results may open new paths for the design of potential vaccines against cutaneous leishmaniasis.

## Results

### γδ T cells are the main source of IL-17A in *Leishmania amazonensis* infection in mice

We have previously demonstrated that *L. amazonensis* infection of Sv129 mice fails to induce expansion of IFN-γ producing CD4 and CD8 T cells, but it triggers the expansion of IL-17A producing γδ T cells ^31^. To determine the main source of IL-17A in *L. amazonensis* infection, we performed a flow cytometry analysis of the draining popliteal lymph node of mice infected in the footpad. *L. amazonensis* infection induced large lesions and a high parasite load in Sv129 mice (Fig.1 a,b). Infected mice showed a higher frequency and number of IL-17A producing T cells than naïve mice (Fig.1 c-e). When we assessed which T cell populations expressed IL-17A in infected mice, we found a large enrichment in γδ T cells in comparison with CD4^+^ or CD8^+^ T cells (Fig.3 f). To analyze whether other lymphocytes could be a source of IL-17A, we performed a multianalysis by flow cytometry. The frequencies of IL-17A-producing CD3^+^CD4^+^ or CD3^+^CD8^+^ T cells did not increase upon infection, however IL-17A-producing CD3^+^γδ^+^ or CD3^+^CD4^-^CD8^-^ γδ^+^ T cells were markedly augmented (Fig. 2 a,c; Supplementary Fig. 1 for gate strategy).

**Fig. 1:**
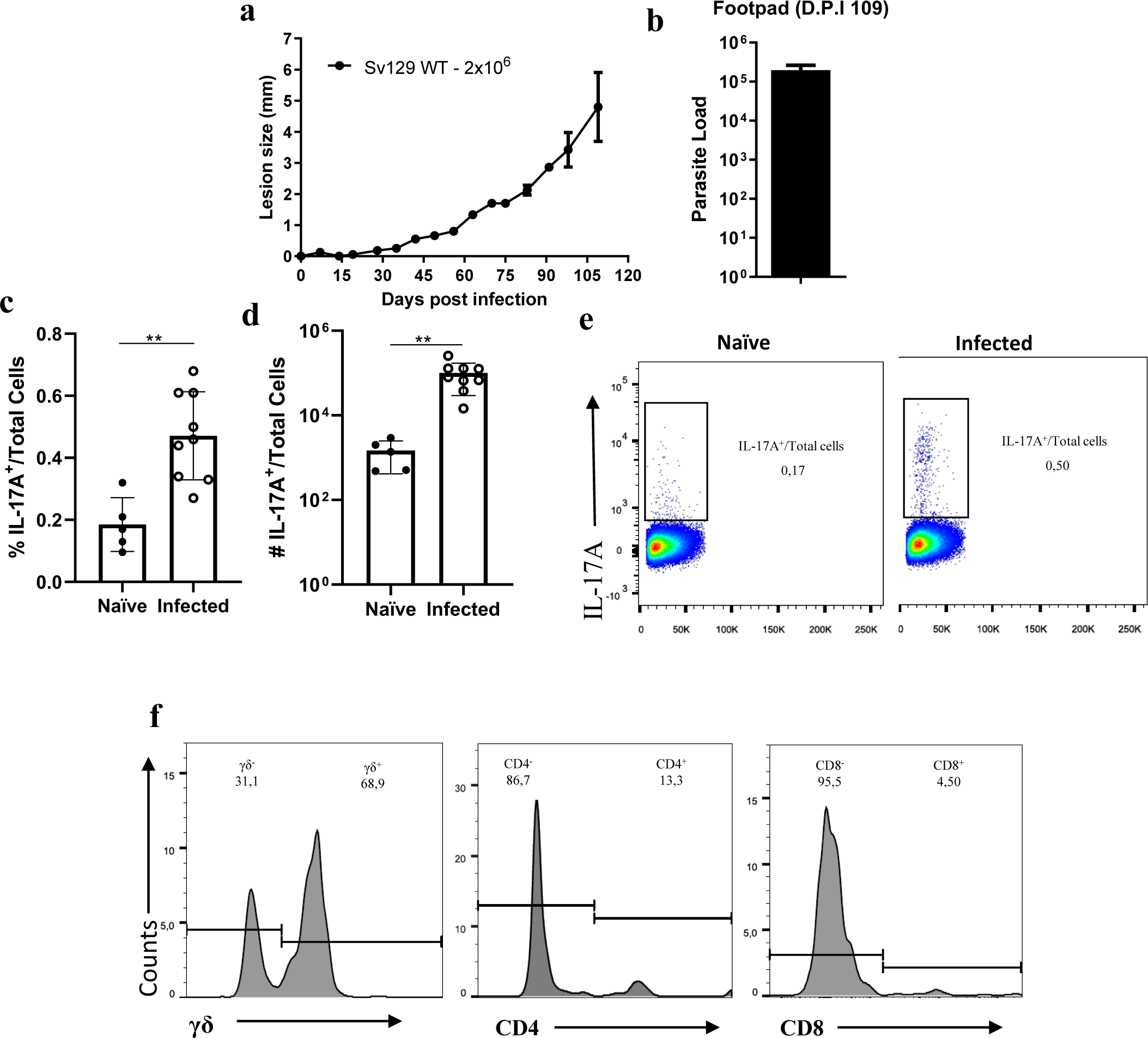
γδ T cells are the major source of IL-17A in *L. amazonensis* infection. Sv129 (WT) mice were infected subcutaneously with 2 x 10^6^ promastigotes of *L. amazonensis* (MHOM/BR/75/Josefa) in the footpad, and the footpad lesion size were followed for 15 weeks. **a** Lesion development; **b** Limiting dilution analysis of parasite burden in the infected footpads; **c, d, e, f** flow cytometry assay of cells of draining popliteal lymph node to quantify the main source of IL-17A. **c** frequency of IL-17A-producing T cells; **d** number of IL-17A-producing T cells; **e** dot plot of frequency of IL-17A-producing T cells; **f** histogram of populations inside of IL-17A-producing T cells. Each dot in the bar graphics represents the value obtained from an individual mouse. Each Dot plot and histogram (**e, g**) represents the mouse value relative to the group average. Data are presented as mean values ± SD. \*\**P* < 0.01, \**P* < 0.05 comparing the indicated groups, as determined by the two-way ANOVA and Student *T* test two-sided. Shown is one representative experiment of three independent experiments performed.

**Fig. 2:**
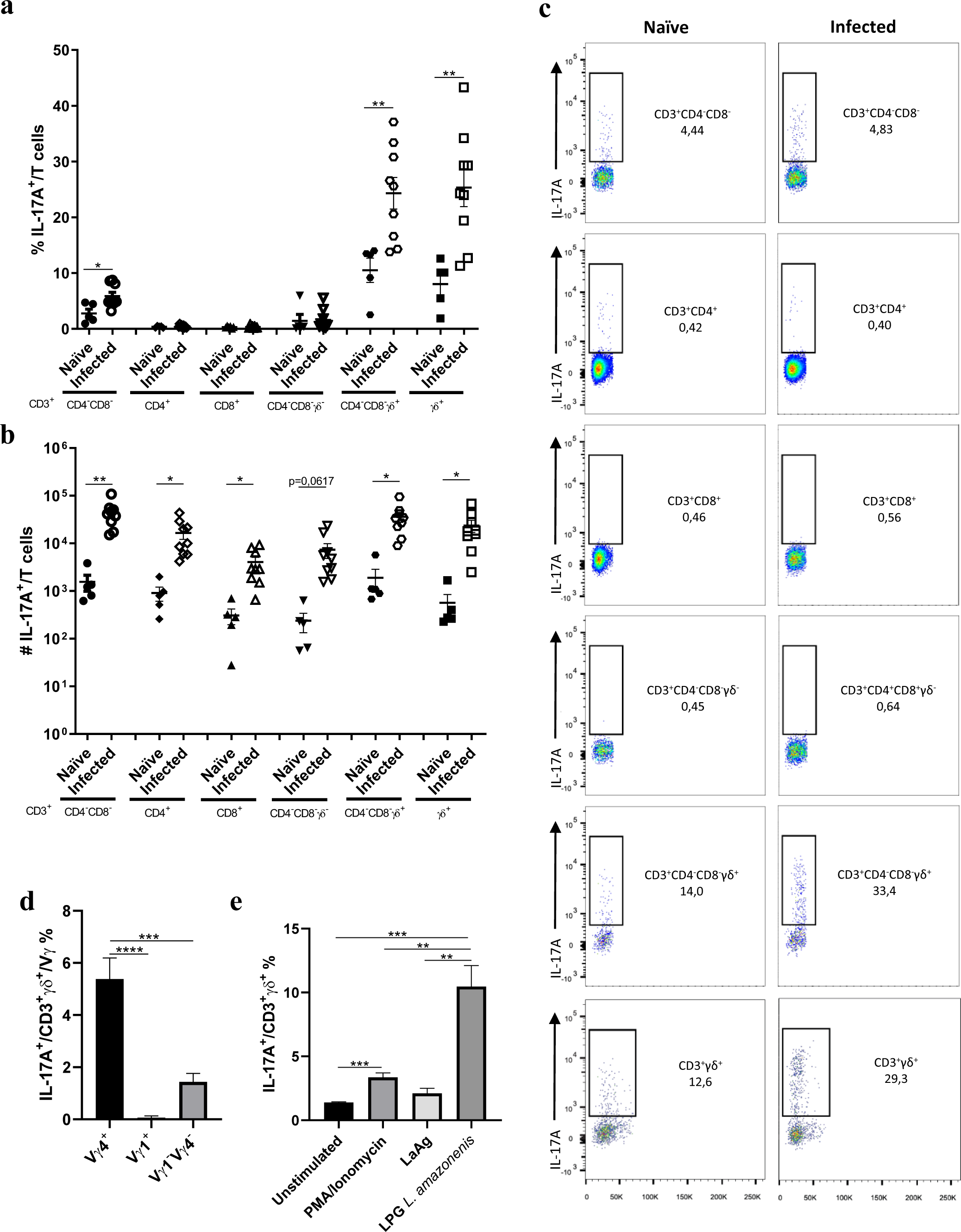
*L. amazonensis* infection induces the accumulation of IL-17A-producing γδ T cells. Sv129 (WT) mice were infected subcutaneously with 2 x 10^6^ promastigotes of *L. amazonensis* (MHOM/BR/75/Josefa) in the footpad, and the footpad lesion size were followed for 15 weeks. **a**, **b** and **c** Flow cytometry analyze of cells of draining popliteal lymph node to measure the main source of IL-17A. **a** frequency of IL-17A-producing T cells; **b** number of IL-17A-producing T cells; **c** dot plot of frequency of IL-17A-producing T cells. **d** frequency of IL-17A-producing γδ T cells stimulated with *L. amazonensis* LPG. Each dot in the graphics represents the value obtained from an individual mouse. Each Dot plot (**c**) represents the mouse value relative to the group average. Data are presented as mean values ± SD. Naïve (n=5), Infected (n=7) \*\**P* < 0.01, \**P* < 0.05 comparing the indicated groups, as determined by the Student *T* test two-sided. Shown is one representative experiment of three independent experiments performed.

**Fig. 3:**
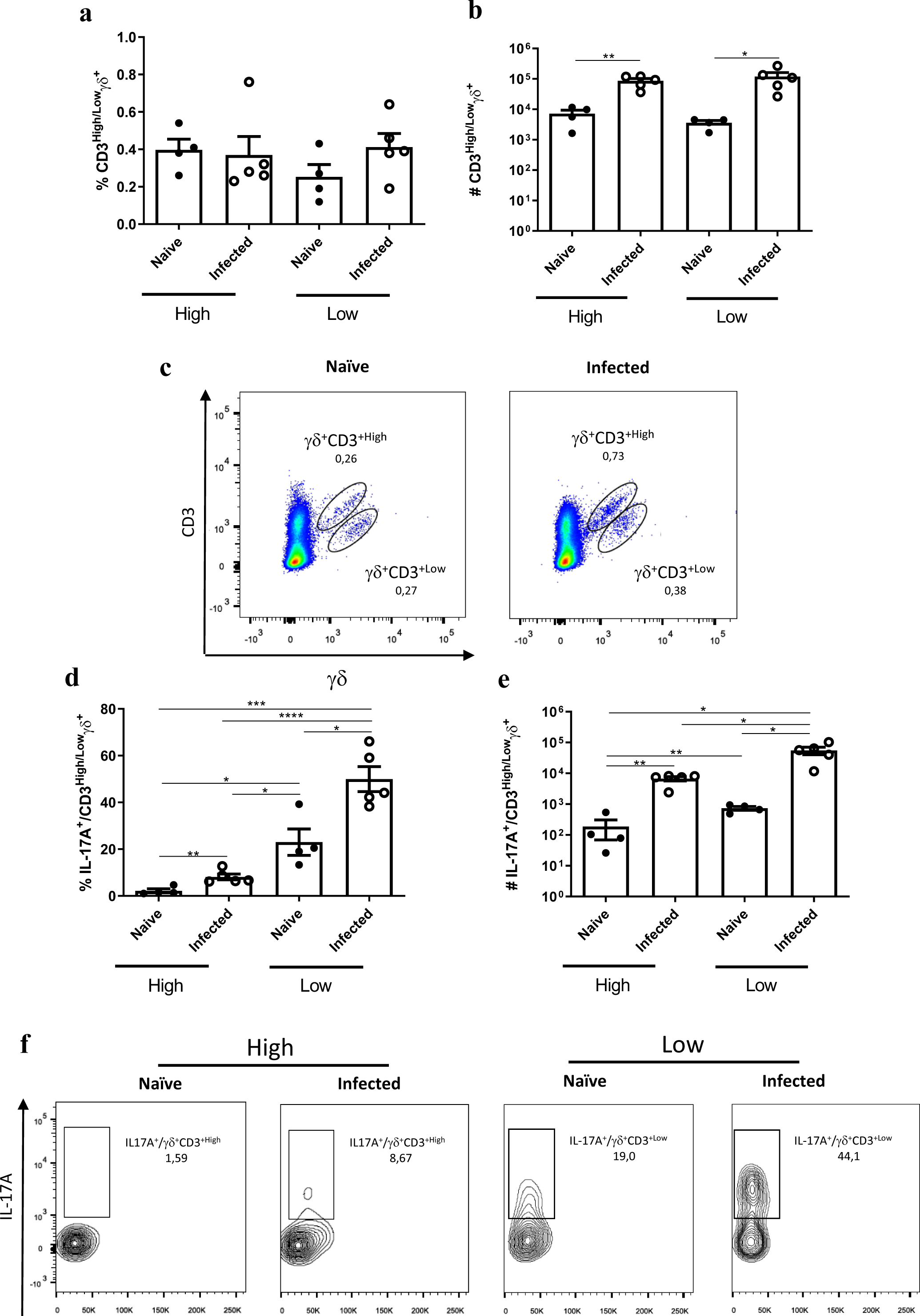
*L. amazonensis* infection induces the preferential expansion of IL-17A-producing CD3^Low^γδ T cells. Sv129 (WT) mice were infected subcutaneously with 2 x 10^6^ promastigotes of *L. amazonensis* (MHOM/BR/75/Josefa) in the footpad, and the footpad lesion size were followed for 15 weeks. **a**, **b** and **c** Flow cytometry analyze of cells of draining popliteal lymph node. **a** frequency of CD3^High^ and CD3^Low^ γδ T cells; **b** number of CD3^High^ and CD3^Low^ γδ T cells; **c** dot plot of frequency of CD3^High^ and CD3^Low^ γδ T cells. Each dot in the graphics represents the value obtained from an individual mouse. Each Dot plot (**c**) represents the mouse value relative to the group average. **d** frequency of IL-17A-producing CD3^Low^γδ T cells; **e** number of IL-17A-producing CD3^Low^γδ T cells; **f** dot plot of frequency of IL-17A-producing CD3^Low^γδ T cells.Data are presented as mean values ± SD. \*\*\*\**P* < 0.0001, \*\*\**P* < 0.001, \*\**P* < 0.01, \**P* < 0.05 comparing the indicated groups, as determined by the Student *T* test two-sided. Shown is one representative experiment of three independent experiments performed.

Given the capacity of γδ T cells to express either CD4^39^ or CD8^40^, we examined the impact of this expression on the induction of IL-17A. Infected mice aren’t able to induce an increase of frequency of IL-17A-producing CD3^+^CD4^+^γδ^+^ T cells, instead of an increase in the number of IL-17A-producing CD3^+^CD4^+^γδ^+^ T cells (Supplementary Fig. 3 a-f), nor in the frequency and number of IL-17A-producing CD3^+^CD8^+^γδ^+^ T cells, instead of an increase in the number of total population of CD3^+^CD8^+^γδ^+^ T cells (Supplementary Fig. 3 g-l). *L. amazonensis* infection induced an increase in the absolute numbers of all IL-17A-producing T cells, except IL-17A-producing CD3^+^CD4^-^CD8^-^γδ^-^ T cells (Fig. 2 b). The increase of number was consistent with the increase of the number of specific and total cells in draining popliteal lymph node of mice infected (Supplementary Fig. 2 a,c). There was no difference in the frequency of specific populations (Supplementary Fig. 2 b). Together, these data reveal that γδ T cells, lacking CD4 and CD8 expression, are the main sources of IL-17A in *L. amazonensis* infection.

Looking at the Vγ usage of the responding γδ T cells, we found that *L. amazonensis* infection induced a higher frequency of IL-17A-producing CD3^+^ γ4^+^δ^+^ T cells (Fig. 2d). As expected, the infection did not induce IL-17A-producing CD3^+^γ1^+^δ^+^ T cells. Additionally, the infection led to a modest increase in the frequency of IL-17A-producing CD3^+^γ4^-^1^-^δ^+^ T cells (Fig. 2d), excluding a substantial contribution in IL-17A production from other populations such as Vγ6^+^ cells. These data collectively suggest that the CD3^+^γ4^+^δ^+^ T cell population constitutes the main source of IL-17A during *L. amazonensis* infection.

To investigate the molecular basis of the γδ T cell response to *L. amazonensis*, we conducted an *in vitro* assay using cells from the draining popliteal lymph node of infected animals, in the presence or absence of LPG, one of the most abundant components of the cell surface of all *Leishmania* species. Interestingly, we observed that stimulation with *L. amazonensis* LPG induced much higher induction of IL-17A-producing γδ T cells compared to stimulation with PMA/Ionomycin or with total lysate of *L. amazonensis* antigens (LaAg) (Fig. 2e). These data suggest that *L. amazonensis* LPG may act as a ligand for γδ T cells and promote their IL-17A response to infection.

### IL-17A-producing γδ T cell response to *L. amazonensis* is not restricted to one strain

To analyze whether other strains of *L. amazonensis* could induce the expansion of IL-17A-producing γδ T cells, we performed a flow cytometry assay of the draining popliteal lymph node of mice infected in the footpad with 2x10^6^ promastigotes of *L. amazonensis* strain MHOM/BR/77/LTB0016. As noted with the strain MHOM/BR/75/Josefa (Fig.1 a,b), the infection by *L. amazonensis* strain MHOM/BR/77/LTB0016 in Sv129 mice induced large lesions and a high parasite load (Supplementary Fig.4 a,b). Infected mice had a higher frequency and number of IL-17A producing T cells than naïve mice (Supplementary Fig.4 c-e). When we looked at which populations were expressing IL-17A in infected mice by strain MHOM/BR/77/LTB0016, we also saw a large enrichment in γδ T cells compared to CD4^+^ or CD8^+^ T cells (Supplementary Fig.4 f). Flow cytometry multianalysis showed that mice infected with strain MHOM/BR/77/LTB0016 failed to induce IL-17A-producing CD3^+^CD4^+^ or CD3^+^CD8^+^ T cells, but accumulated IL-17A-producing CD3^+^γδ^+^ and CD3^+^CD4^-^CD8^-^γδ^+^ T cells (Supplementary Fig.5 a,c; Supplementary Fig. 1 for gating strategy).

Infected mice by strain MHOM/BR/77/LTB0016 also aren’t able to induce an increase of frequency of IL-17A-producing CD3^+^CD4^+^γδ^+^ T cells, instead of an increase in the number of IL-17A-producing CD3^+^CD4^+^γδ^+^ T cells (Supplementary Fig.6 a-f). The infection induced a small increase in the frequency and number IL-17A-producing CD3^+^CD8^+^γδ^+^ T cells, instead of do not induce an increase in the frequency and number of total populations of CD3^+^CD8^+^γδ^+^ T cells (Supplementary Fig.6 g-l). We found an increase in the number of total cells in draining popliteal lymph node of mice infected (Supplementary Fig. 7a), but not in CD3^+^CD4^+^ or CD3^+^CD4^-^CD8^-^γδ^-^ T cells(Supplementary Fig. 7 b,c). These data demonstrate that the dominant IL-17A-producing γδ T cell response to *L. amazonensis* infection is not restricted to one strain of the parasite, but is likely a conserved feature of CL.

### IL-17A-producing CD3^Low^γδ T cells are expanded in infected Sv129 mice

In our analyses, we observed that γδ T cells were distributed in two cell groups, expressing high (CD3^High^) or low (CD3^Low^) levels of the CD3 complex (Supplementary Fig. 1 for gating strategy). We therefore decided to investigate whether CD3 levels could segregate with the expression of IL-17A by γδ T cells in infected Sv129 mice. Although there was no difference in the frequency of CD3^High^ and CD3^Low^ γδ T cells between infected and naïve mice (Fig. 3 a,c), their absolute numbers increased in relation to naïve mice (Fig. 3b). Interestingly, infected Sv129 mice had a higher percentage (Fig. 3 d,f) and absolute number (Fig. 3 e) of IL-17A-producing CD3^Low^γδ T cells compared to CD3^High^γδ T cells from infected mice. We also found that this expansion of IL-17A-producing CD3^Low^γδ T cells was similarly induced by the strain MHOM/BR/77/LTB0016 (Supplementary Fig. 8). Thus, the IL-17A response to *L. amazonensis* infection in Sv129 mice is mostly confined to CD3^Low^γδ T cells.

### Type 1 IFNR signaling controls the expansion of IL-17A-producing CD3^Low^ γδ T cells and limits disease severity

Given the importance of type 1 IFNR signaling in controlling the expansion of γδ T cells^41^, we assessed its impact on the dynamics of the γδ T cell response to *L. amazonensis* infection. We evaluated the profile of development of the lesion size between Sv129 WT and A129 mice (deficient in type 1 IFNR) and observed that A129 mice were more susceptible to infection by *L. amazonensis*, presenting larger lesion sizes (Fig. 4 a,b) and higher parasite loads (Fig. 4c). Next, we analyzed the impact of type 1 IFNR signaling on γδ T cells. When we gated both populations of γδ T cells there was no difference in the frequency and absolute number of total γδ T cells (Supplementary Fig. 9 a-c) nor IL-17A-producing γδ T cells (Supplementary Fig. 9 d-f). The lack of type 1 IFNR signaling had a dual effect on reducing the fraction of CD3^Low^ γδ T cells but increasing their production of IL-17A when compared to the same populations from Sv129 WT mice or to CD3^High^ γδ T cells from A129 mice (Fig. 4 f,h,i). However, we did not observe a difference in the absolute number of any population analyzed (Fig. 4 e,g), nor was there any distinction in the cellularity of the draining popliteal lymph node of infected animals (Supplementary Fig. 9h). Together, these data demonstrate that type 1 IFNR signaling may preferentially contain the expansion of IL-17A-producing CD3^Low^ γδ T cells while controlling *L. amazonensis* infection and its pathological manifestation.

**Fig. 4:**
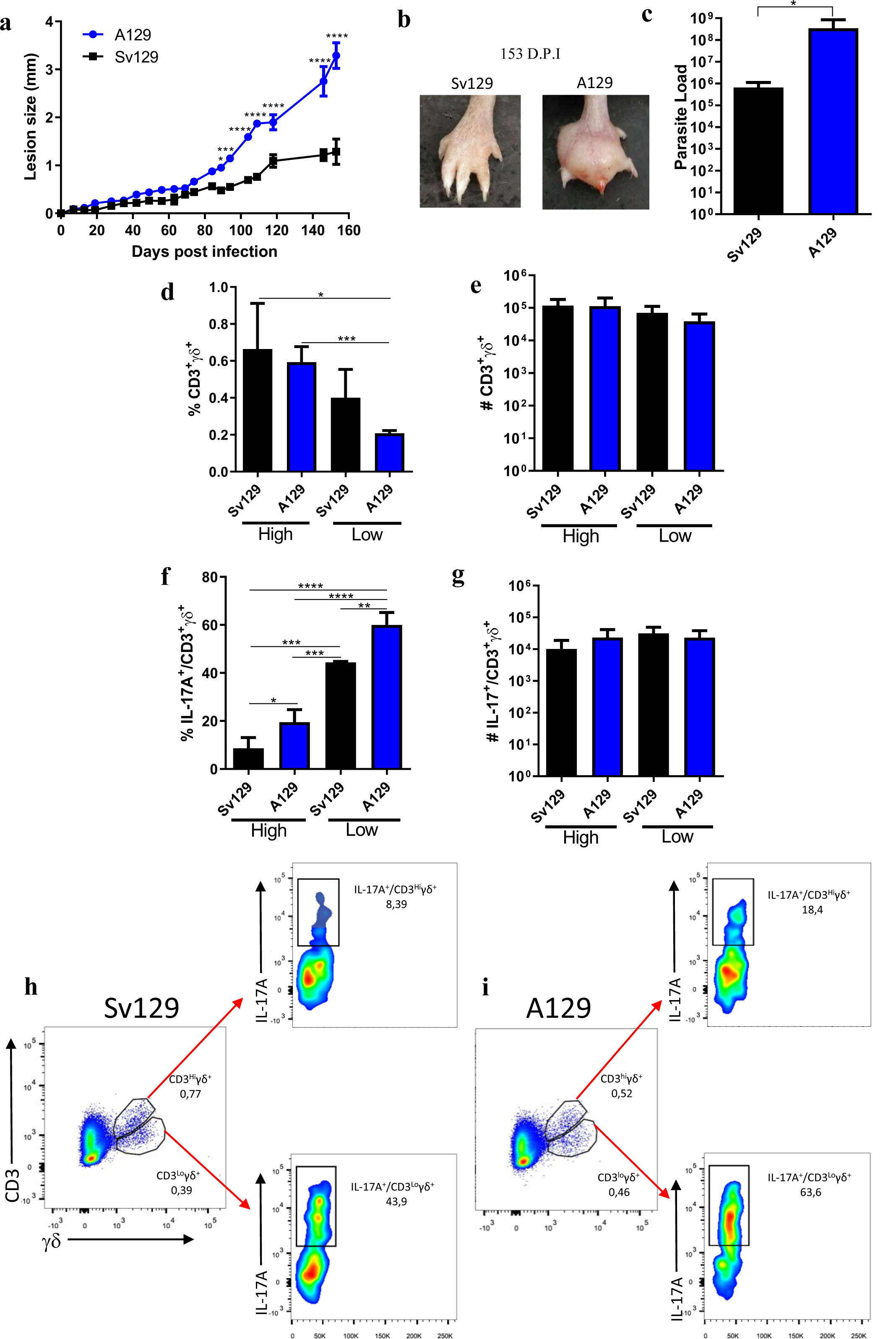
IL-17A-producing CD3^Low^ γδ T cells expand in the absence of type 1 IFNR in *L. amazonensis* infection. Sv129 (WT) and A129 mice were infected subcutaneously with 2 x 10^6^ promastigotes of *L. amazonensis* (MHOM/BR/75/Josefa) in the footpad, and the footpad lesion size were followed for 21 weeks. **a** Lesion development; **b** images of footpads infected **c** Limiting dilution analysis of parasite burden in the infected footpads; **d, e, f, g, h and i** flow cytometry assay of cells of draining popliteal lymph node. **d** frequency of CD3^High^ γδ T cells and CD3^Low^ γδ T cells; **e** number of CD3^High^ γδ T cells and CD3^Low^ γδ T cells; **f** frequency of IL-17A-producing CD3^Low^ γδ T cells; **g** number of IL-17A-producing CD3^Low^ γδ T cells; **h** dot plot of frequency of CD3^High^ γδ T cells, CD3^Low^ γδ T cells and IL-17A-producing CD3^Low^ γδ T cells in Sv129 (WT) mice; **i** dot plot of frequency of CD3^High^ γδ T cells, CD3^Low^ γδ T cells and IL-17A-producing CD3^Low^ γδ T cells in A129 mice. Each Dot plot (**h, i**) represents the mouse value relative to the group average. Data are presented as mean values ± SD. Sv129 n=3, A129= 4 \*\**P* < 0.01, \**P* < 0.05 comparing the indicated groups, as determined by the two-way ANOVA, Student *T* test two-sided and Mann-Whitney. Shown is one representative experiment of two independent experiments performed.

### γδ T cells-IL-17-elastase axis is pathogenic in *L. amazonensis* infection

To understand the role of γδ T cells and associate pathways in *L. amazonensis* infection, we infected TCRδ^-/-^ mice that lack all γδ T cells, IL-17RA^-/-^ that lack the capacity to respond to IL-17, and NE^-/-^ that lack the signaling of neutrophil elastase. During the first 50 days post-infection there was no difference in the lesion between TCRδ^-/-^ or IL-17RA^-/-^ compared to WT mice (Fig. 5a). However, whereas WT mice continued to grow the lesion after that time point, reaching the peak phase around 70 days d.p.i., both TCRδ^-^/^-^ and IL-17RA^-^/^-^ mice were able to control the lesion (Fig. 5a). Next, when we compared NE-deficient and WT mice, we observed a similar profile (Fig. 5b). These data suggest that the pathogenesis of *L. amazonensis* infection involves γδ T cells, signaling through the IL17 receptor and neutrophil elastase, likely in an IL-17A-mediated axis.

**Fig. 5:**
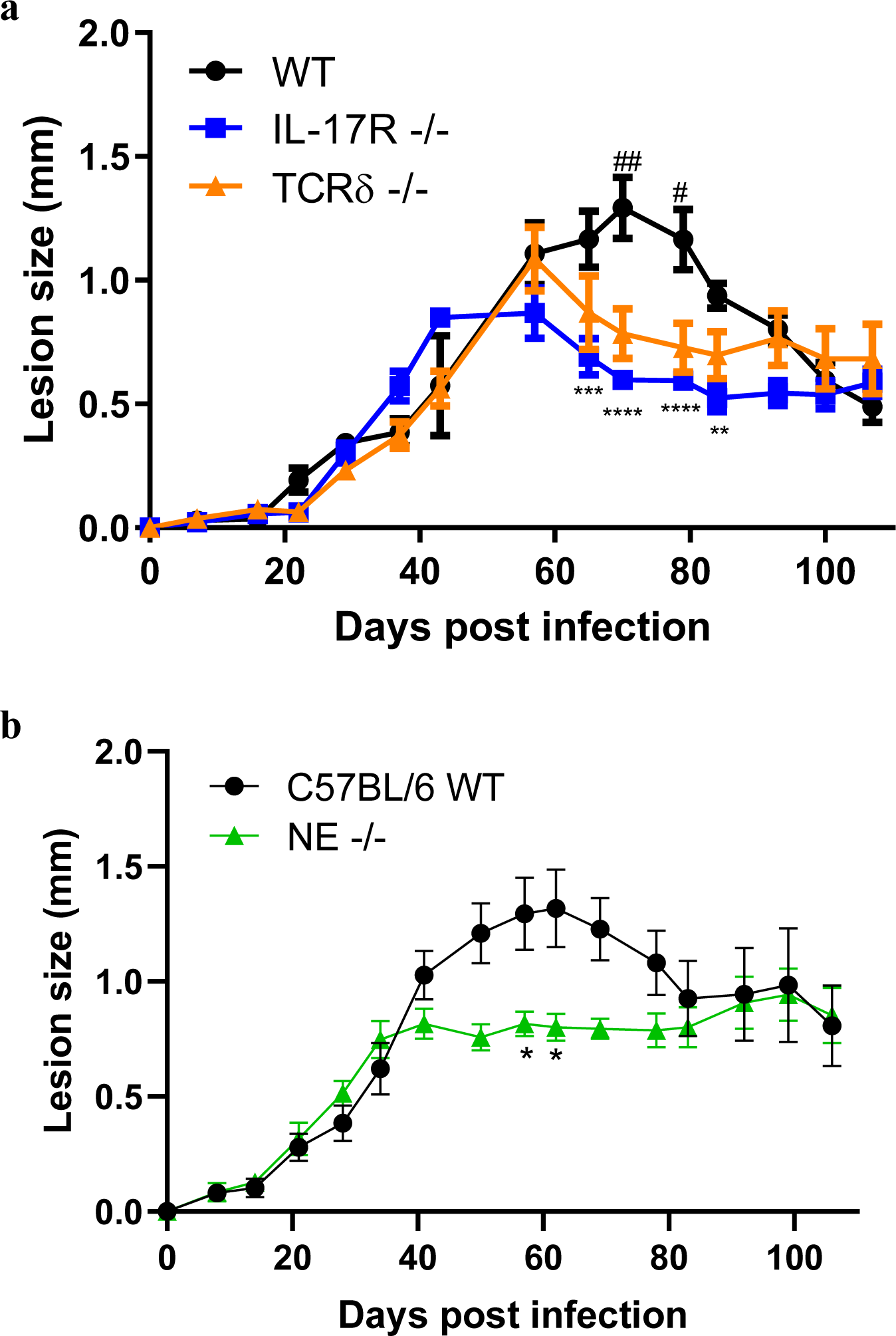
γδ T cells, IL-17 and neutrophil elastase are important to control *L. amazonensis* infection. C57BL/6 (WT), TCRδ^-/-^, IL-17RA^-/-^, and NE^-/-^ mice were infected subcutaneously with 2 x 10^6^ promastigotes of *L. amazonensis* (MHOM/BR/75/Josefa) in the footpad were followed for 15 weeks. **a** Lesion development of WT (n=5), TCRδ^-/-^ (n=5) and IL-17RA^-/-^ (n=5), the # symbol represent the difference WT and TCRδ^-/-^, * symbol represent the difference WT and IL-17RA^-/-^; **b** Lesion development of WT and NE^-/-^. Data are presented as mean values ± SD. \*\*\*\**P* < 0.0001,\*\**P* < 0.01, \**P* < 0.05 comparing the indicated groups, as determined by the two-way ANOVA. Shown is one representative experiment of three independent experiments performed.

### Transfer of IFNγ-biased γδ T cells protect against *L. amazonensis* infection

To determinate the impact of an adoptive transfer of γδ T cells during *L. amazonensis* infection in Sv129 mice, we sorted γδ^+^ T cells and γδ^-^ T cells and we transferred into naïve mice, one hour before the footpad infection of 2x10^6^ *L. amazonensis* promastigotes. Strikingly, we observed that mice that received γδ^+^ T cells were able to control the development of the progressive lesion, in contrast with mice receiving γδ^-^ T cells or PBS as control (Fig. 6a). We also tested the impact of doubling the dose of transferred γδ^+^ T cells, and observed better control of lesion compared to control mice that received PBS (Fig. 6b). Of note, the transfer of both amounts of cells led to similar reductions in the parasite load (Fig 6c). These data demonstrate the strong potential of γδ T cell adoptive transfer to control cutaneous leishmaniasis caused by *L. amazonensis*.

**Fig. 6:**
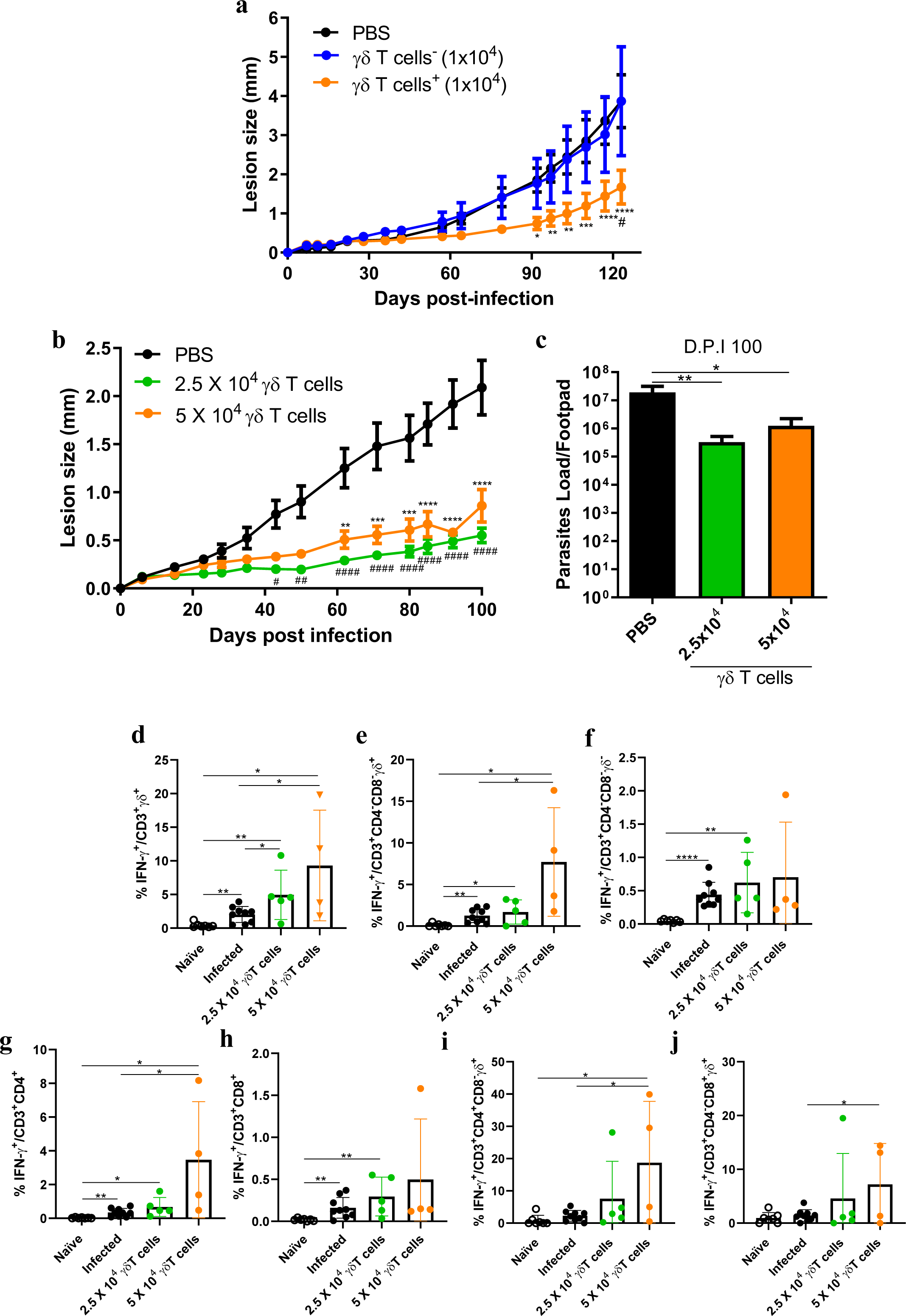
Adoptive transfer of IFNγ-prone γδ T cells protect Sv129 mice against *L. amazonensis* infection. Sv129 naïve mice received the transfer of cells (20 draining popliteal lymph nodes from mice were pooled to obtained γδ T cells) and 1 hour later were infected subcutaneously with 2 x 10^6^ promastigotes of *L. amazonensis* (MHOM/BR/75/Josefa) in the footpad and were followed for 17 weeks **a** and 14 weeks in **b**. **a** Lesion development of PBS (n=5), γδ^-^ T cells (n=5) and γδ^+^ T cells (n=5)-received mice. * symbol represent the difference PBS and γδ^+^ T cells -received mice, # symbol represent the difference γδ^-^ T cells and γδ^+^ T cells -received mice **b** Lesion development of PBS (n=5), 2.5x10^4^ (n=5) and 5x10^4^ (n=4) γδ^+^ T cells -received mice. * symbol represent the difference PBS and 2.5x10^4^ γδ^+^ T cells -received mice, # PBS and 5x10^4^ γδ^+^ T cells -received mice **c** Limiting dilution analysis of parasite burden in the infected footpads. The frequency of IFN-γ-producing **d** CD3^+^γδ^+^ **e** CD3^+^CD4^-^CD8^-^γδ^+^ **f** CD3^+^CD4^-^ CD8^-^γδ^-^ **g** CD3^+^CD4^+^ **h** CD3^+^CD8^+^ **i** CD3^+^CD4^+^CD8^-^γδ^+^ and **j** CD3^+^CD4^-^CD8^+^γδ^+^ T cells. Data are presented as mean values ± SD. \*\*\*\**P* < 0.0001, \*\*\**P* < 0.001, \*\**P* < 0.01, \**P* < 0.05 comparing the indicated groups, as determined by the two-way ANOVA, Student *T* test two-sided and Mann-Whitney test. Shown is two representative experiments of five experiments performed.

To understand how the adoptive transfer of γδ^+^ T cells could provide a protective response against *L. amazonensis* infection, we analyzed the draining popliteal lymph nodes of the infected mice at 14 weeks post-transfer. We observed that the transfer of 2.5 and 5x10^4^ γδ^+^ T cells induced a higher frequency of IFN-γ-producing CD3^+^γδ^+^ T cells in comparison to infected and naïve mice (Fig. 6d). The amount of 2.5 γδ^+^ T cells was not able to induce the increase in frequency in comparison to infected mice, just was more than naïve mice. However, 5x10^4^ γδ^+^ T cells were able to induce higher frequency of IFN-γ-producing CD3^+^CD4^-^CD8^-^γδ^+^ in comparison to infected and naïve mice (Fig. 6e). The transfer of 5x10^4^ γδ^+^ T cells were capable to induce a higher frequency of IFN-γ-producing CD3^+^CD4^+^CD8^-^γδ^+^ (Fig. 6i) the was more than infected and naïve mice, and also induced an increase in CD3^+^CD4^-^CD8^+^γδ^+^ T cells that was more than naïve mice (Fig. 6J). These data show that the FACS-sorted total γδ T cells that provide protection against *L. amazonensis* infection upon adoptive transfer into Sv129 mice trigger an IFNγ response.

We also analyzed the cytotoxic potential of the adoptively transferred γδ T cells. *L. amazonensis* infection did not induce granzyme B, perforin-producing and LAMP-1^+^-expressing CD3**^+^**γδ^+^ T cells, however, the transfer of 2.5 or 5x10^4^ γδ^+^ T cells induced a higher frequency LAMP-1^+^-expressing CD3**^+^**γδ^+^ T cells (Supplementary Fig. 10a). The transfer of γδ^+^ T cells did not impact in granzyme B and perforin-producing CD3**^+^**γδ^+^ T cells expansion (Supplementary Fig. 10 b,c). The transfer of 2.5 or 5x10^4^ γδ^+^ T cells induced an increase in the frequency of LAMP-1^+^-expressing CD3^+^CD4^+^ T cells in comparison to naïve mice (Supplementary Fig. 10d). In addition, the transfer of 2.5x10^4^ γδ^+^ T cells increase the expression of LAMP-1^+^-expressing CD3^+^CD8^+^ T cells in comparison to infected and naïve mice. Together, these result show that the transfer of γδ^+^ T cells induced a higher frequency of cytotoxic-related-CD107a.

### CD3^Low^ γδ T cells control the development of the lesion in *L. amazonensis* infection

To analyze the role of CD3^High^ and CD3^Low^ γδ T cells during *L. amazonensis* infection, we sorted these cells and transferred them for naïve mice. One hour later, we infected these mice with 2x10^6^ promastigotes of *L. amazonensis* in the footpad. We evaluated that PBS and CD3^High^ γδ T cells-received mice had the same profile of development of the lesion (Supplementary Fig. 11a). Around 116 days, CD3^Low^ γδ T cells began to control the development of the progressive lesion in comparison to PBS-received mice. Later, around 137 days, CD3^Low^ γδ T cells began to control the development of the progressive lesion in comparison to CD3^High^ γδ T cells-received mice (Supplementary Fig. 11a). However, we did not observe the difference in parasite load (Supplementary Fig. 11b). These data indicate that CD3^Low^ γδ T cells can reduce the lesion of *L. amazonensis* infection.

### γδ T cell-mediated protection is widespread to other strains of mice

To evaluate the impact of γδ T cell transfer in other mouse models, we infected BALB/c and C57BL/6 mice with 2x10^6^ promastigotes of *L. amazonensis* in the footpad. First, we observed that γδ T cell transfer controlled lesion in genetically susceptible BALB/c mice compared to mice receiving only PBS that began to develop progressive lesion (Fig. 7a). Then, we analyzed that γδ T cell transfer controlled the lesion size in genetically resistant C57BL/6 mice, during an earlier phase of infection, until the peak phase, compared to animals that received only PBS, which managed to contain the development of the lesion (Fig. 7b). These results firmly demonstrate the potential of γδ T cells to protect different species of mice against *L. amazonensis* infection.

**Fig. 7:**
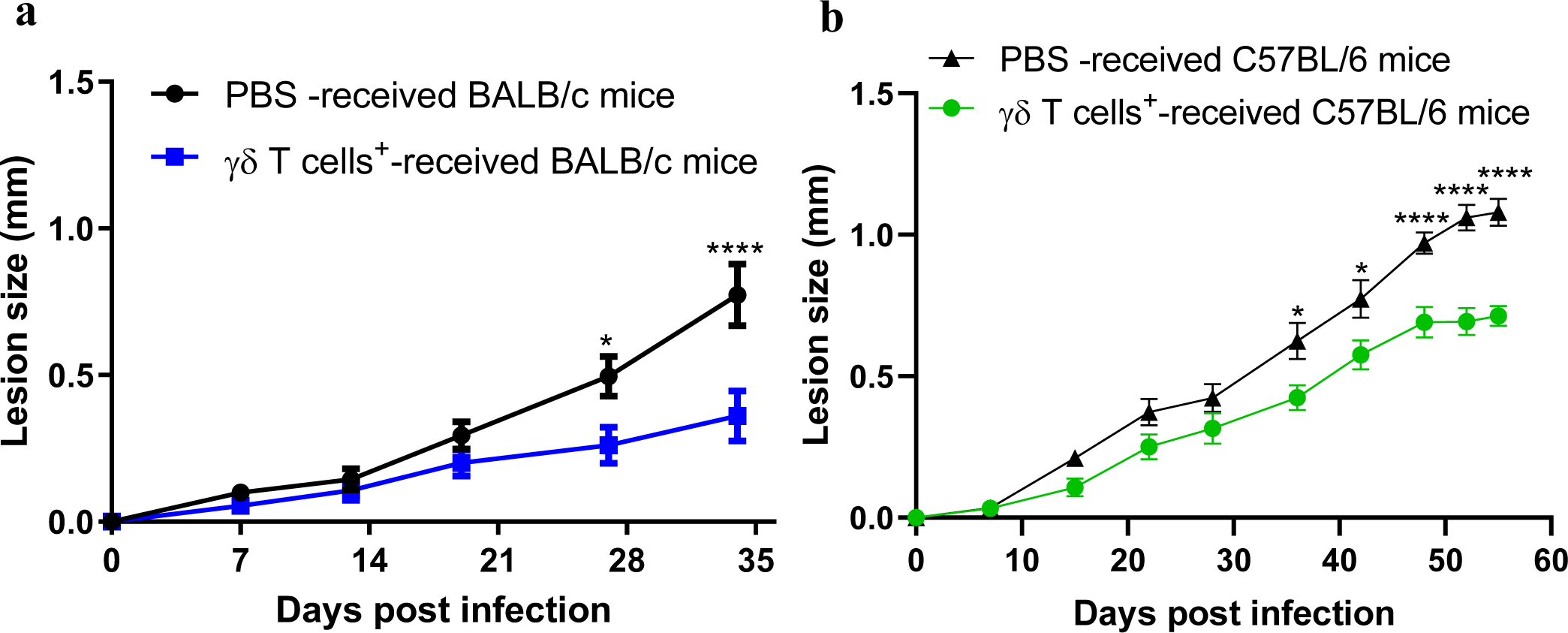
Adoptive transfer of γδ T cells protect BALB/c and C57BL/6 mice against *L. amazonensis* infection. BALB/c and C57BL/6 naïve mice received the transfer of 2.5x10^4^ γδ T cells (20 draining popliteal lymph nodes from mice of each strain were pooled to obtained γδ T cells) and 1 hour later were infected subcutaneously with 2 x 10^6^ promastigotes of *L. amazonensis* (MHOM/BR/75/Josefa) in the footpad and were followed for 5 weeks in BALB/c mice and 8 weeks in C57BL/6 mice. **a** Lesion development of PBS (n=9) and γδ T cells (n=10)-received BALB/c mice **b** Lesion development of PBS (n=8) and γδ T cells(n=8)-received C57BL/6 mice. Data are presented as mean values ± SD. \*\*\*\**P* < 0.0001,\*\**P* < 0.01, \**P* < 0.05 comparing the indicated groups, as determined by the Student *T* test two-sided. Shown is one representative experiment of twoexperiment performed.

## Discussion

For the first time, this work brings to light the functional dichotomy of γδ T cells in experimental infection by *L. amazonensis*. We found that γδ T cells, in particular their Vγ4^+^ subpopulation, are the main source of IL-17A upon *L. amazonensis* infection in Sv129 mice. Furthermore, we discovered that LPG induces the proliferation of IL-17A-producing γδ T cells. We revealed that the type 1 IFNR signaling is essential to control *L. amazonensis* infection and limits the expansion of IL-17A-producing γδ T cells. We demonstrated that the γδ T cell/IL-17/elastase axis is associated with the pathogenic response. By contrast, when we sorted total γδ T cells and transferred them to mice, we observed control of *L. amazonensis* infection, which associated with an increase in the frequency of IFN-γ-producing γδ T cells. Thus, we propose that the functional dichotomy between IL-17A and IFN-γ production by γδ T cells impacts distinct outcomes of experimental infection by *L. amazonensis*.

For a long time, the presence of γδ T cells has been found in lesions of *L. braziliensis* infected-patients with cutaneous and mucocutaneous leishmaniasis^30^, as well as enriched in human peripheral blood mononuclear cells (PBMCs) of *L. major* infected-patients^27^. Although γδ T cells were initially associated with the early stages of granuloma formation of *L. braziliensis* infection^26^, these cells were present in variable amounts in the CL, mucosal (ML) and disseminated cutaneous leishmaniasis (DisCL) lesions, regardless of the clinical characteristics (and duration) of the lesions^28^.

Our previous study demonstrated that γδ T cells expand in *L.amazonensis*-infected Sv129 mice, and using γδ T cell-deficient mice, we demonstrated that these cells are pathogenic during *L. amazonensis* infection ^31^. By contrast, γδ T cell deficiency did not impact in the lesion size and parasite load in mice infected with *L. major*^42^ or *L. guyanensis*^43^. This difference could be explained by the fact that *L. amazonensis* infection is capable of inducing an anergic state in CD4 and CD8 T cells through the induction of PD-1 expression, consequently leading to an incapacity to produce IFN-γ^17^. This scenario could favor the expansion of γδ T cells during *L. amazonensis* infection, given that PD-1 expression is associated with the cytotoxic role and IFN-γ producing by these cells^44^.

However, treatment with anti-TCRδ antibody led to higher lesion size and parasite load during *L. major* infection in BALB/c mice, and delayed the healing of cutaneous lesion in CBA/J model^45^. The same effect was observed when the antibody was administered to mice following oral immunization with LaAg^38^.

We previously showed that *L.amazonensis* infection induces an increase of IL-17A-producing γδ T cells in Sv129 mice ^31^. However, their precise contribution to the IL-17A response in *L. amazonensis* infection was not known before the present study. IL-17 has been associated with the severity of *L. braziliensis*^46,47^, *L. major*^48^, *L. tropica*^49^ and *L. panamensis*^50^ infections. Higher levels of IL-17 were found in ML lesions than CL, and in PBMCs from patients in relation to healthy individuals^46–49^. Furthermore, there is higher expression of IL-17A^+^ cells in lesions of patients with DCL, compared to localized cutaneous leishmaniasis (LCL), as well as to healthy individuals and LCL from *L.braziliensis*-infected patients^23^. It was previously observed that in lesions of *L. braziliensis*-infected patients, CD4^+47,48^, CD8^+47^ and CD14^+47^ cells can be sources of IL-17. In experimental cutaneous leishmaniasis, in *L. major*-infected BALB/c mice, the main source of IL-17A were CD4^+^ T cells and, to a lesser extent, CD8^+^ and γδ T cells^21^. However, in C57BL/6 mice, the main source of IL-17 were γδ T cells, with a secondary contribution of CD4^+^ and CD8^+^ T cells^22^. In this study, we demonstrated that Vγ4^+^ γδ T cells are the major source of IL-17A in infection with different strains of *L. amazonensis*.

A key function of IL-17A is to recruit ^21,50^neutrophils to the site of infection ^21,51^, where they have been associated with facilitating macrophage infection as a reservoir for *Leishmania*^52,53^. In mouse strains exhibiting resistance, neutrophils are promptly recruited within hours following parasite inoculation, but their presence diminishes to 1%–2% by day 3 post-infection. Conversely, in susceptible BALB/c mice, neutrophils continue to be recruited and persist in substantial numbers at the infection site for more than 10 days following parasite inoculation^54^. Furthermore, Th17 cells, IL-17 and neutrophils increase up to 90 days after *L. mexicana* infection. This suggests that chronic inflammation during infection is a consequence of IL-17-mediated neutrophil recruitment^55^. As a potential effector mechanism of neutrophils, here we explored neutrophil elastase (NE), a serine protease normally expressed in neutrophil primary granules and are implicated in regulating the formation of neutrophil extracellular traps (NETs)^56^. *L. amazonensis* can induce the release of these NETs by neutrophils^57^. Furthermore, IL-17 and NETs were colocalized at infection sites in the chronic phase in *L. mexicana infection*, indicating that this IL-17/NETs axis may contribute to sustaining inflammation^55^. Our data suggest that the absence of NE has a similar pathology phenotype in *L. amazonensis* infection as a deficient γδ T cell/IL-17 axis, namely a lower lesion size in the respective mouse models.

Supporting our proposed γδ T cell/IL-17/NE axis, neutrophils have been shown to release elastase upon co-culture with TCR-activated γδ T cells^58^. In addition, NE added to γδ T-cell cultures in the absence of neutrophils, reproduced the increase in the activation of γδ T cells observed in co-cultures of γδ T cells and neutrophils^58^. When pre-incubated with a specific NE inhibitor, the effect in of T cell activation was abolished^58^. However, this profile depends on the activation status of the γδ T cells, as neutrophils were previously reported to inhibit (E)-4-hydroxy-3-methylbut-2-en-1-yl diphosphate (HMB-PP)-induced activation of the major subset of human γδ T cells, Vγ9Vδ2 T cells^59^. It is worth noting that the inhibition of human γδ T cells activated by HMB-PP only occurs with viable neutrophils, co-culture with apoptotic neutrophils leads to increased activation of γδ T cells activated by HMBPP^59^.

γδ T cell activation would be the key event to trigger the proposed γδ T cell/IL-17/NE axis in our infection model. Most γδ T cells recognize non-peptidic molecules independently of the major histocompatibility complex (MHC)^60^. Lipophosphoglycans (LPGs), the main glycoconjugates identified on the surface membranes of *Leishmania* parasites, are consistently expressed in promastigote forms. In the case of *L. amazonensis*, depending on the strain, LPGs display polymorphisms in their structures and can differentially immunomodulate NO and cytokine production by murine macrophages^61,62^ Additionally, LPG serves as a virulence factor, playing a direct role in the attachment and establishment of parasites within the midgut of sand flies^63,64^. Furthermore, Leishmania’s LPG can induce exhaustion molecules such as PD-1 in lymphocytes and their ligands PD-L2 in macrophages, potentially contributing to infection^65,66^. Importantly, we found that *L. amazonensis*’ LPG can induce the expansion of IL-17A-producing γδ T cells. These data suggest that the LPG of *L. amazonensis* may serve as a γδ T cell ligand and could contribute to the pathogenesis of the infection.

We observed that naïve Sv129 mice possess two populations of γδ T cells and that *L. amazonensis* infection induces an increase in IL-17A-producing CD3^Low^ γδ T cells compared to uninfected mice. This data reveals an important characteristic of γδ T cells in *L. amazonensis* infection, since CD3^High^ γδ T cells had been previously characterized as the largest producers of IL-17A^67,68^. Subsequent experiments will be necessary to determine which mechanisms regulate IL-17A-producing CD3^Low^ cells in *L. amazonensis* infection. Our data are in line with the findings that in naïve BALB/c and C57BL/6 mice that γδ T cells can be subdivided into two populations according to CD3 expression (CD3^High^ and CD3^Low^)^67,68^.

A129 mice (deficient in type 1 IFNR) were more susceptible to infection by *L. amazonensis* strain Josefa, presenting higher lesion size and parasite load. However, it is possible that this profile is dependent on the *Leishmania* strain and background of mice, as observed the opposite profile in infection with *L. amazonensis* strain LTB0016^69^. In addition, in experimental *L. major* infection in C57BL/6 and BALB/c mice, there is no difference in lesion size and parasite burden between IFN-β and type 1 IFNR-deficient mice compared to WT mice^70^. We identified that the lack of type 1 IFNR signaling did not impact the frequency and number of total γδ T cells, as well as those IL-17A-producing γδ T cells, but increased the percentage of IL-17A-producing CD3^Low^ γδ T cells increased compared to that of IL-17A-producing CD3^High^ γδ T cells. These data suggest a possible pathway for the immunopathogenesis of cutaneous leishmaniasis caused by *L. amazonensis* strain Josefa via the expansion of IL-17A-producing CD3^Low^ γδ T cells, which is favored in the absence of type 1 IFNR signaling.

Finally, we explored the therapeutic potential of the adoptive transfer of IFNγ-biased total γδ T cells. It has been shown that CD27^+^ IFNγ-prone γδ T cells expand substantially more than CD27^−^ IL-17-biased γδ T cells upon adoptive transfer; and the stimulation via the TCR greatly favors IFNγ over IL-17A producers^71,72^). Previously, γδ T cell transfer was shown to be capable of controlling *Trypanosoma* infection seemingly through IFN-γ-producing γδ T cells^73^. A similar protection is observed in *Toxoplasma* infection^74^. For the first time, we demonstrated that γδ T cell transfer can protect against *L. amazonensis* infection in different mouse models. The protection was associated with an increase in the frequency of IFN-γ-producing γδ T cells. This finding is consistent with previous data on the GL3 antibody stimulating γδ T cells to produce IFN-γ^75^. Interestingly, blockade of IFNγ increases IL-17 level, neutrophils and, consequently, the pathology in *L.major* infection^51^, supporting the hypothesis that, in C57BL/6 mice, IFNγ may be critical for negatively regulating the IL-17/neutrophil responses.

Furthermore, it has been described that CD3^+^CD4^-^CD8^-^ T cells are the main cells que express CD107a in lesions of patients with *L. braziliensis* infection^76^. ^57^Our results demonstrate that *L. amazonensis* infection induced a higher frequency of CD107a-expressing CD3^+^γδ^+^ T cells. Although we did not observe any difference in the expression of granzyme B and perforin by γδ^+^ T cells, more studies are needed to evaluate the participation of granzyme A and K^77^. Mainly because it has already been demonstrated that granzyme A induces cell death in the absence of granzyme b and perforin^78^. Therefore, the increase in IFN-γ and cytotoxicity may contribute to controlling the disease and not be associated with pathogenicity.

Altogether, our data demonstrate for the first time the impact of the functional dichotomy of γδ T cells on cutaneous leishmaniasis: whereas *L. amazonensis* infection naturally activates IL-17A-producing γδ T cells that promote disease pathogenesis, the adoptive transfer of γδ T cells that trigger an IFN-γ response *in vivo* is able to control the pathogenesis of *L. amazonensis* infection in mice. As human γδ T cells are strongly (more than mice) biased towards IFN-γ (instead of IL-17A) production^79,80^, we believe our findings may have important therapeutic implications.

## Acknowledgements

We would like to thank Dr. Natacha G. Sousa, Dr. Alda Cruz and Dr. professor Bartira Rossi-Bergmann for scientific disucssion. We received financial support from Cientista do Nosso Estado (Faperj – E-26/200.993/2022); Productivity Fellowships from Conselho Nacional de Desenvolvimento Científico e Tecnológico (307632/2022-9) and Agency for Support and Evaluation of Graduate Education (CAPES) Finance code 001.

## Methods

### Animals

Female Sv129 WT^31^ and deficient in type 1 IFNR (A129)^81^ mice in the Sv129 background were obtained from the Laboratory of Inflammation and Immunity at the Paulo de Góes Microbiology Institute of the Federal University of Rio de Janeiro (UFRJ). Strains deficient in TCRδ (TCRδ^-^/^-^), IL-17 Receptor (IL-17RA ^-^/^-^) and elastase neutrophil (NE^-^/^-^) in the C57BL/6 background were obtained from the Ribeirão Preto Medical School of the University of São Paulo (FMRP/USP). All animals were between 6 and 8 weeks old. Mice were maintained at mice facility of the Paulo de Góes Microbiology Institute in ventilated cages in an ambient temperature and humidity-controlled room with a 12 h light/12 h dark cycle with continuous access to food and water. All experiments with animals were previously approved by the Ethics Committee with the Use of Animals in Scientific Experimentation (CEUA) of the Health Sciences Center of UFRJ, under protocol number A07-21-037-20.

### Parasites,infection and LPG purification

Two strains of *Leishmania (Leishmania) amazonensis* originally isolated from cutaneous leishmaniasis were used in this work: MHOM/BR/75/Josefa^82^ and MHOM/BR/77/LTB0016^83^. LPG from *L. amazonensis* (Josefa strain) were extracted and purified as previously reported^61^. Parasites were routinely isolated from lesions of infected BALB/c mice and cultured in M199 medium (Sigma-Aldrich, Darmstadt, He, Germany) supplemented with 0.02% hemin, 10% heat-inactivated fetal bovine serum (or FBS; HIFCS, GIBCO Laboratories, Grand Island, NY, USA), 100 U/L penicillin and 100 μg/L, and maintained at 26 °C. For the infection assays, stationary phase parasites (from cultures used up to the 5^th^ passage) were washed with PBS three times (1000 x g, 10 min). The cell pellet was then resuspended in PBS for *in vivo* infection. Animals were infected in the right hind footpad (subcutaneously), using a syringe (HAMILTON) with 2 × 10^6^ promastigotes of both strains of *L. amazonensis* in 20 μL for the characterization of the infection. The volume of 10 μL (intradermal) was used by infection then sorting experiments. The lesion size was monitored using a Mitutoyo caliper.

### Limiting Dilution Assay (LDA)

To determinate the parasite load, LDA was performed. Briefly, infected footpads were removed and macerated in 1 mL of M199 medium (Sigma-Aldrich, Darmstadt, He, Germany) containing 10% FBS. A 96-well plate was pre-filled with 150 μl M199 medium (Sigma-Aldrich, Darmstadt, He, Germany) supplemented with 10% FBS (HIFCS, GIBCO Laboratories, Grand Island, NY, USA), and 50 μL of the cell suspension was placed in the first well, and a 1:4 serial dilution was performed by passing 50 μL of the dilution to the following well for a total of 24 dilutions for each sample. The plates were incubated in a bio-oxygen demand (BOD) incubator at 26°C for 7 to 14 days. After this time, the last well to show growth of promastigotes, as observed visually on a light microscope, was used to calculate the total number of parasites present in the organ. The calculation used was as follows: Number of parasites = 4^x^ (where x is the number of the last well in which parasites were observed).

### Flow Cytometry

Cells from draining popliteal lymph nodes (1 x 10^6^) were stimulated for 4 h at 37 °C with PMA (phorbol 12-myristate 13-acetate, 10 ng/mL, Sigma-Aldrich, Darmstadt, He, Germany) and Ionomycin (10 ng/mL, Sigma-Aldrich, Darmstadt, He, Germany), LPG of *L. amazonensis* (strain Joseja, 10 ug/mL) or LaAg (5 ug/mL) in the presence of a Golgi complex inhibitor Brefeldin A (5 mg/mL, Biolegend, SanDiego, CA, USA). All centrifugation steps were performed at 4 °C. The cells were washed with PBS by centrifugation at 400 g for 5 min at 4°C, blocked with 5 µL/well Anti-Mo CD16/CD32 (clone 93) (eBiosicence, Carlsbad, CA, USA) (1:100) for 15 min, followed by 5 µL/well of the antibody cocktail for extracellular staining: anti-CD3-BV421 (clone 17A2) (BioLegend, SanDiego, CA, USA) (1:200); anti-CD4-APC-CY-7 (clone GK1.5) (BioLegend, SanDiego, CA, USA) (1:200); anti-CD4-FITC (clone GK1.5) (eBioscience, Carlsbad, CA, USA) (1:200); anti-CD8-Percp-Cy-5.5 (clone 53-6.7) (BioLegend, SanDiego, CA, USA) (1:200); anti-TCRδ-BV510 (Clone GL3) (BioLegend, SanDiego, CA, USA) (1:200); anti-TCRδ-FITC (Clone GL3) (eBioscience, Carlsbad, CA, USA); anti-TCR Vγ4-FITC (Clone UC310A6) (eBioscience, Carlsbad, CA, EUA); anti-TCR Vγ1.1-Percp-Cy-5.5 (Clone 2.11) (eBioscience, Carlsbad, CA, EUA) (1:200) (1:200); anti-CD107a-PE-Cy-7 (clone 1D4B) (BioLegend, SanDiego, CA, USA) (1:200) and incubation for 30 min at 4°C in the dark. Cells were washed with a cytometry buffer solution (PBS with 5% FBS) then fixed and permeabilized (FoxP3 permeabilization/fixation kit; eBiosicence, Carlsbad, CA, USA) according to the manufacturer’s protocol. The intracellular staining was performed using anti-IL-17A-APC (Clone TC11-18H10.1) (BioLegend, SanDiego, CA, USA) (1:100); anti-IFN-γ-PE (Clone XMG1.2) (BioLegend, SanDiego, CA, USA) (1:100); anti-perforin-PE (Clone S16009A) (BioLegend, SanDiego, CA, USA) (1:100); anti-granzyme B-Alexa Fluor 647 (Clone GB11) (BioLegend, SanDiego, CA, USA) (1:100) for 1 hour at 4°C in the dark. At the end, Cells were washed, resuspended in cytometry buffer solution, and stored in the dark at 4°C until acquisition. Analysis was performed in the FlowJo software vX.0.7 and the graphics were made in GraphPad Prism 8.0.2.

### Cell sorting

Cells from draining popliteal lymph nodes were collected from mice with chronic (Sv129 and C57BL/6) and progressive BALB/c lesions. Then the cells were labeled as described above using antibodies anti-CD3-BV421 (clone 17A2) (BioLegend, SanDiego, CA, USA) (1:200) and anti-TCRδ-BV510 (Clone GL3) (BioLegend, SanDiego, CA, USA) (1:200). Cells were separated using FACSARIA III Cell Sorter and Diva software. Naïve mice received 100µl of PBS (control) or 100µl of PBS containing specific amounts of cells in the venous sinus by retro-orbital injection as previously described^84^.

### Statistical analysis

Analysis of the results was performed using the GraphPad Prism 8.0.2program. Statistical tests of significance for differences between groups of mice were determined by the Two-way ANOVA test for the analysis of lesion development, by the Student’s T test and Mann-Whitney test when there were only two groups to be compared. Values are represented as mean ± standard deviation of the mean.

## Supplementary figure legends

**Supplementary Fig. 1: Gating strategy to identify the main source of IL-17 in *L. amazonensis* infection.** Sv129 (WT) mice were infected subcutaneously with 2 x 10^6^ promastigotes of *L. amazonensis* (MHOM/BR/75/Josefa) in the footpad, and the footpad lesion size were followed for 18 weeks. The gate strategy represents flow cytometry assay of cells of draining popliteal lymph node to quantify the main source of IL-17A. CD3^+^ γδ^+^ T cells and IL-17A producing T cells were gated from single cells. The histogram of γδ^+^, CD4^+^ and CD8^+^ T cells were gated from IL-17A producing T cells. IL-17A-producing CD3^+^ CD4^+^, CD3^+^ CD8^+^, CD3^+^ CD4^-^ CD8^-^, CD3^+^ CD4^+^ γδ ^+^ and CD3^+^ CD8^+^ γδ ^+^ T cells were gated from CD3^+^. IL-17A-producing CD3^+^ CD4^-^ CD8^-^γδ^+^ and CD3^+^ CD4^-^ CD8^-^ γδ^-^ T cells were gated from CD3^+^ CD4^-^ CD8^-^. Each Dot plot represents the mouse value relative to the group average. Shown is one representative experiment of three independent experiments performed.

**Supplementary Fig. 2: *L. amazonensis* infection induces an increase in cell numbers in draining popliteal lymph node.** Sv129 (WT) mice were infected subcutaneously with 2 x 10^6^ promastigotes of *L. amazonensis* (MHOM/BR/75/Josefa) in the footpad, and the footpad lesion size were followed for 18 weeks. **a**, **b** and **c** Flow cytometry analyze of cells of draining popliteal lymph node. **a** number of total cells; **b** frequency of T cells; **c** number of T cells. Each dot in the graphics represents the value obtained from an individual mouse. Data are presented as mean values ± SD. Naïve (n=5), Infected (n=7) \*\*\**P* < 0.001, \*\**P* < 0.01, \**P* < 0.05 comparing the indicated groups, as determined by the Student *T* test two-sided. Shown is one representative experiment of three independent experiments performed.

**Supplementary Fig. 3: *L. amazonensis* infection fails to induce an expansion of IL-17A-producing CD3^+^CD4^+^γδ^+^ and CD3^+^CD8^+^γδ^+^ T cells.** Sv129 (WT) mice were infected subcutaneously with 2 x 10^6^ promastigotes of *L. amazonensis* (MHOM/BR/75/Josefa) in the footpad, and the footpad lesion size were followed for 15 weeks. **a**, **b**, **c**, **d**, **e**, **f**, **g**, **h**, **i**, **j**, **k** and **l** Flow cytometry analyze of cells of draining popliteal lymph node to measure the main source of IL-17A. **a** frequency of CD3^+^CD4^+^γδ^+^ T cells; **b** number of CD3^+^CD4^+^γδ^+^ T cells; **c** dot plot of frequency of CD3^+^CD4^+^γδ^+^ T cells; **d** frequency of IL-17A-producing CD3^+^CD4^+^γδ^+^ T cells; **e** number of IL-17A-producing CD3^+^CD4^+^γδ^+^ T cells; **f** contour plot of frequency of IL-17A-producing CD3^+^CD4^+^γδ^+^ T cells; **g** frequency of CD3^+^CD8^+^γδ^+^ T cells; **h** number of CD3^+^CD8^+^γδ^+^ T cells; **i** dot plot of frequency of CD3^+^CD8^+^γδ^+^ T cells; **j** frequency of IL-17A-producing CD3^+^CD8^+^γδ^+^ T cells; **k** number of IL-17A-producing CD3^+^CD8^+^γδ^+^ T cells; **l** contour plot of frequency of IL-17A-producing CD3^+^CD8^+^γδ^+^ T cells; Each dot in the graphics represents the value obtained from an individual mouse. Each dot and contour plot (**c, f, I, l**) represents the mouse value relative to the group average. Data are presented as mean values ± SD. Naïve (n=5), Infected (n=7) \*\**P* < 0.01, \**P* < 0.05 comparing the indicated groups, as determined by the Student *T* test two-sided. Shown is one representative experiment of three independent experiments performed.

**Supplementary Fig. 4: IL-17A-producing T cells induced by *L. amazonensis* (strain MHOM/BR/77/LTB0016) infection are also enriched in γδ T cells.** Sv129 (WT) mice were infected subcutaneously with 2 x 10^6^ promastigotes of *L. amazonensis* (MHOM/BR/77/LTB0016) in the footpad, and the footpad lesion size were followed for 18 weeks. **a** Lesion development; **b** Limiting dilution analysis of parasite burden in the infected footpads; **c, d, e, f** flow cytometry assay of cells of draining popliteal lymph node to quantify the main source of IL-17A. **c** frequency of IL-17A-producing T cells; **d** number of IL-17A-producing T cells; **e** dot plot of frequency of IL-17A-producing T cells; **f** histogram of populations inside of IL-17A-producing T cells. Each dot in the bar graphics represents the value obtained from an individual mouse. Each Dot plot and histogram (**e, g**) represents the mouse value relative to the group average. Data are presented as mean values ± SD. Naïve (n=5), Infected (n=5) \*\**P* < 0.01 comparing the indicated groups, as determined by the Student *T* test two-sided. Shown is one representative experiment of two independent experiments performed.

**Supplementary Fig. 5: *L. amazonensis* (strain MHOM/BR/77/LTB0016) infection induces a higher expansion of IL-17A-producing γδ T cells.** Sv129 (WT) mice were infected subcutaneously with 2 x 10^6^ promastigotes of *L. amazonensis* (MHOM/BR/77/LTB0016) in the footpad, and the footpad lesion size were followed for 18 weeks. **a**, **b** and **c** Flow cytometry analyze of cells of draining popliteal lymph node to measure the main source of IL-17A. **a** frequency of IL-17A-producing T cells; **b** number of IL-17A-producing T cells; **c** dot plot of frequency of IL-17A-producing T cells. Each dot in the graphics represents the value obtained from an individual mouse. Each Dot plot (**c**) represents the mouse value relative to the group average. Data are presented as mean values ± SD. Naïve (n=5), Infected (n=5) \*\**P* < 0.01, \**P* < 0.05 comparing the indicated groups, as determined by the Student *T* test two-sided. Shown is one representative experiment of two independent experiments performed.

**Supplementary Fig. 6: *L. amazonensis* (strain MHOM/BR/77/LTB0016) infection fails to induce the expansion of IL-17A-producing CD3^+^CD4^+^γδ^+^ and CD3^+^CD8^+^γδ^+^ T cells.** Sv129 (WT) mice were infected subcutaneously with 2 x 10^6^ promastigotes of *L. amazonensis* (MHOM/BR/77/LTB0016) in the footpad, and the footpad lesion size were followed for 18 weeks. **a**, **b**, **c**, **d**, **e**, **f**, **g**, **h**, **i**, **j**, **k** and **l** Flow cytometry analyze of cells of draining popliteal lymph node to measure the main source of IL-17A. **a** frequency of CD3^+^CD4^+^γδ^+^ T cells; **b** number of CD3^+^CD4^+^γδ^+^ T cells; **c** dot plot of frequency of CD3^+^CD4^+^γδ^+^ T cells; **d** frequency of IL-17A-producing CD3^+^CD4^+^γδ^+^ T cells; **e** number of IL-17A-producing CD3^+^CD4^+^γδ^+^ T cells; **f** contour plot of frequency of IL-17A-producing CD3^+^CD4^+^γδ^+^ T cells; **g** frequency of CD3^+^CD8^+^γδ^+^ T cells; **h** number of CD3^+^CD8^+^γδ^+^ T cells; **i** dot plot of frequency of CD3^+^CD8^+^γδ^+^ T cells; **j** frequency of IL-17A-producing CD3^+^CD8^+^γδ^+^ T cells; **k** number of IL-17A-producing CD3^+^CD8^+^γδ^+^ T cells; **l** contour plot of frequency of IL-17A-producing CD3^+^CD8^+^γδ^+^ T cells; Each dot in the graphics represents the value obtained from an individual mouse. Each dot and contour plot (**c, f, I, l**) represents the mouse value relative to the group average. Data are presented as mean values ± SD. Naïve (n=5), Infected (n=5) \*\**P* < 0.01, \**P* < 0.05 comparing the indicated groups, as determined by the Student *T* test two-sided. Shown is one representative experiment of two independent experiments performed.

**Supplementary Fig. 7: *L. amazonensis* (strain MHOM/BR/77/LTB0016) infection induces an increase in cell numbers in draining popliteal lymph node.** Sv129 (WT) mice were infected subcutaneously with 2 x 10^6^ promastigotes of *L. amazonensis* (MHOM/BR/77/LTB0016) in the footpad, and the footpad lesion size were followed for 18 weeks. **a**, **b** and **c** Flow cytometry analyze of cells of draining popliteal lymph node. **a** number of total cells; **b** frequency of T cells; **c** number of T cells. Each dot in the graphics represents the value obtained from an individual mouse. Data are presented as mean values ± SD. Naïve (n=5), Infected (n=5) \*\**P* < 0.01, \**P* < 0.05 comparing the indicated groups, as determined by the Student *T* test two-sided. Shown is one representative experiment of two independent experiments performed.

**Supplementary Fig. 8: *L. amazonensis* (strain MHOM/BR/77/LTB0016) infection induces higher expansion of IL-17A-producing CD3^Low^γδ T cells.** Sv129 (WT) mice were infected subcutaneously with 2 x 10^6^ promastigotes of *L. amazonensis* (MHOM/BR/77/LTB0016) in the footpad, and the footpad lesion size were followed for 18 weeks. **a-f** Flow cytometry analyze of cells of draining popliteal lymph node. **a** frequency of CD3High and CD3Low γδ T cells; **b** number of CD3High and CD3Low γδ T cells; **c** dot plot of frequency of CD3High and CD3Low γδ T cells. Each dot in the graphics represents the value obtained from an individual mouse. Each Dot plot (**c**) represents the mouse value relative to the group average. **d** frequency of IL-17A-producing CD3^Low^γδ T cells; **e** number of IL-17A-producing CD3^Low^γδ T cells; **f** dot plot of frequency of IL-17A-producing CD3^Low^γδ T cells. Each dot in the graphics represents the value obtained from an individual mouse. Each dot plot (**f**) represents the mouse value relative to the group average. Data are presented as mean values ± SD. Naïve (n=5), Infected (n=5) \*\*\*\**P* < 0.0001, \*\*\**P* < 0.001, \*\**P* < 0.01, \**P* < 0.05 comparing the indicated groups, as determined by the Student *T* test two-sided. Shown is one representative experiment of two independent experiments performed.

**Supplementary Fig. 9: Type 1 IFNR is important to control the expansion of IL-17A-producing CD3^Low^ γδ T cells in *L. amazonensis* infection.** Sv129 (WT) and A129 mice were infected subcutaneously with 2 x 10^6^ promastigotes of *L. amazonensis* (MHOM/BR/75/Josefa) in the footpad, and the footpad lesion size were followed for 21 weeks. **a, b, c, d, e, f and g** flow cytometry assay of cells of draining popliteal lymph node. **a** frequency γδ T cells; **b** number of γδ T cells; **c** dot plot of frequency of γδ T cells **d** frequency of IL-17A-producing γδ T cells; **e** number of IL-17A-producing γδ T cells; **f** dot plot of frequency of IL-17A-producing γδ T cells; **g** number of total T cells. Each Dot plot (**c, f**) represents the mouse value relative to the group average. Data are presented as mean values ± SD. Sv129 n=3, A129 n=4. Shown is one representative experiment of two independent experiments performed.

**Supplementary Fig. 10: Transfer of γδ^+^ T cells leads to an expansion of CD107a^+^-producing γδ T cells during *L. amazonensis* infection.** Sv129 naïve mice received the transfer of cells (20 draining popliteal lymph nodes from mice were pooled to obtained γδ T cells) and 1 hour later were infected subcutaneously with 2 x 10^6^ promastigotes of *L. amazonensis* (MHOM/BR/75/Josefa) in the footpad and were followed for 14 weeks. The frequency of CD107a^+^-producing **a** CD3^+^γδ^+^ **b** CD3^+^CD4^-^ CD8^-^γδ^+^ **c** CD3^+^CD4^-^CD8^-^γδ^-^ **d** CD3^+^CD4^+^ and **e** CD3^+^CD8^+^ T cells. Data are presented as mean values ± SD. \*\**P* < 0.01, \**P* < 0.05 comparing the indicated groups, as determined by Student *T* test two-sided. Shown is one representative experiment of two experiments performed.

**Supplementary Fig. 11: CD3^Low^ γδ T cells protect Sv129 mice against *L. amazonensis* infection.** Sv129 naïve mice received the transfer of cells (10 draining popliteal lymph nodes from mice of each strain were pooled to obtained γδ T cells) and 1 hour later were infected subcutaneously with 2 x 10^6^ promastigotes of *L. amazonensis* (MHOM/BR/75/Josefa) in the footpad and were followed for 21 weeks. **a** Lesion development of PBS (n=6), 6x10^4^ CD3^High^ γδ T cells (n=5) and 1.89x10^5^ CD3^Low^ γδ T cells (n=5)-received mice. * symbol represent the difference between PBS and CD3^Low^ γδ T cells -received mice, # symbol represent the difference CD3^High^ γδ T cells and CD3^Low^ γδ T cells -received mice **b** Limiting dilution analysis of parasite burden in the infected footpads. Data are presented as mean values ± SD. \*\*\*\**P* < 0.0001, \*\**P* < 0.01, \**P* < 0.05 comparing the indicated groups, as determined by the two-way ANOVA and Student *T* test two-sided. Shown is one representative experiment of one experiment performed.

