## Supplemental Figures for "IL-17A/IFN-γ producing γδ T cell functional dichotomy impacts cutaneous leishmaniasis in mice"

Sup Fig1

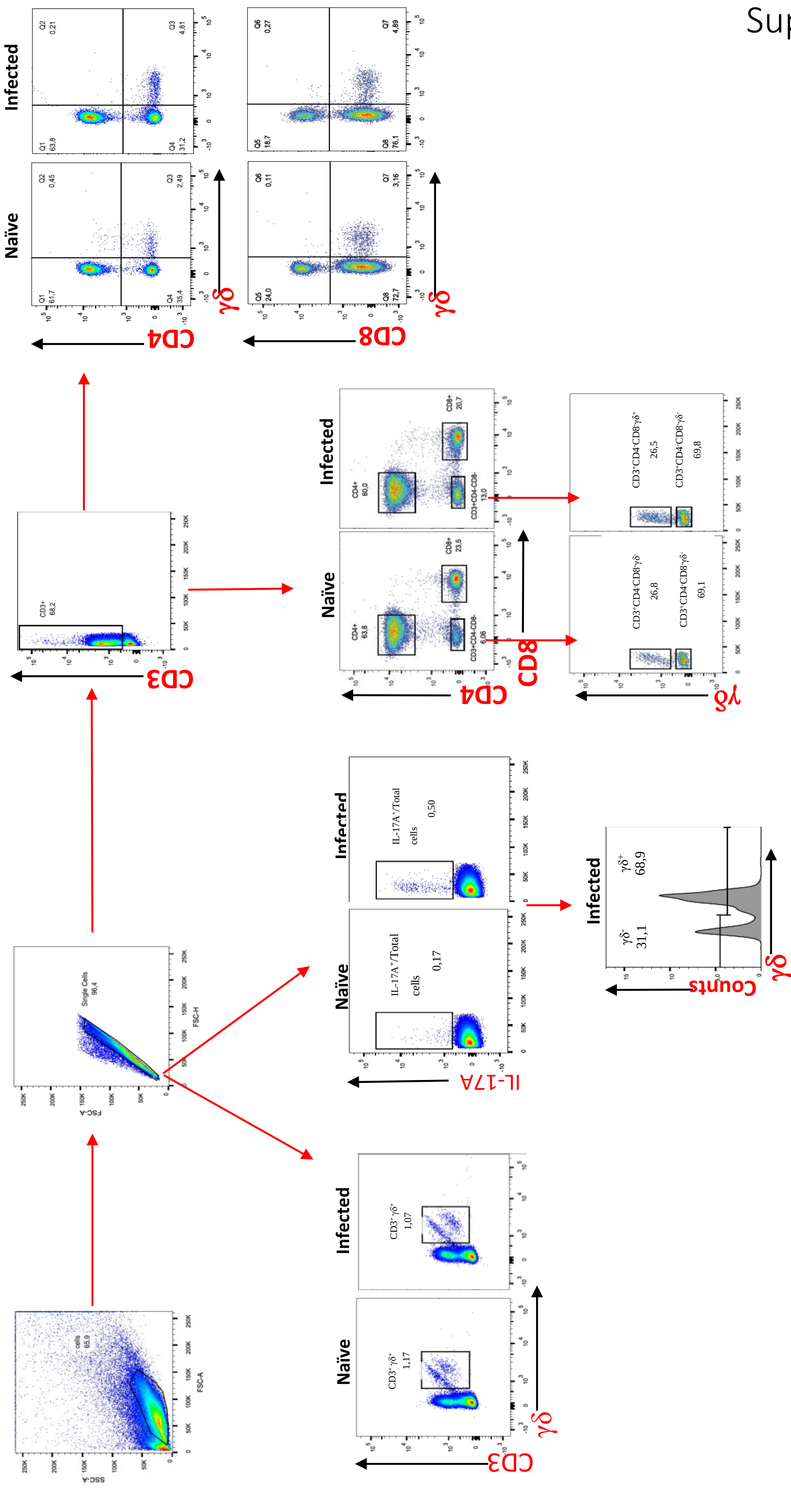

Naïve
Infected
cells
65,9
Single Cells
96,4
CD3+
68,2
CD3
CD4
CD8
γδ
IL-17A
Counts
CD3⁺γδ⁺
1,17
1,07
IL-17A⁺/Total cells
0,17
0,50
γδ⁻
31,1
γδ⁺
68,9
CD4+
63,8
60,0
CD8+
23,5
20,7
CD3+CD4-CD8-
6,06
13,0
CD3⁺CD4⁻CD8⁻γδ⁻
26,8
CD3⁺CD4⁻CD8⁻γδ⁻
69,1
CD3⁺CD4⁻CD8⁻γδ⁺
26,5
CD3⁺CD4⁻CD8⁻γδ⁻
69,8
Q1
61,7
Q2
0,45
Q3
2,49
Q4
35,4
Q1
63,8
Q2
0,21
Q3
4,81
Q4
31,2
Q5
24,0
Q6
0,11
Q7
3,16
Q8
72,7
Q5
18,7
Q6
0,27
Q7
4,89
Q8
76,1

Sup Fig 2

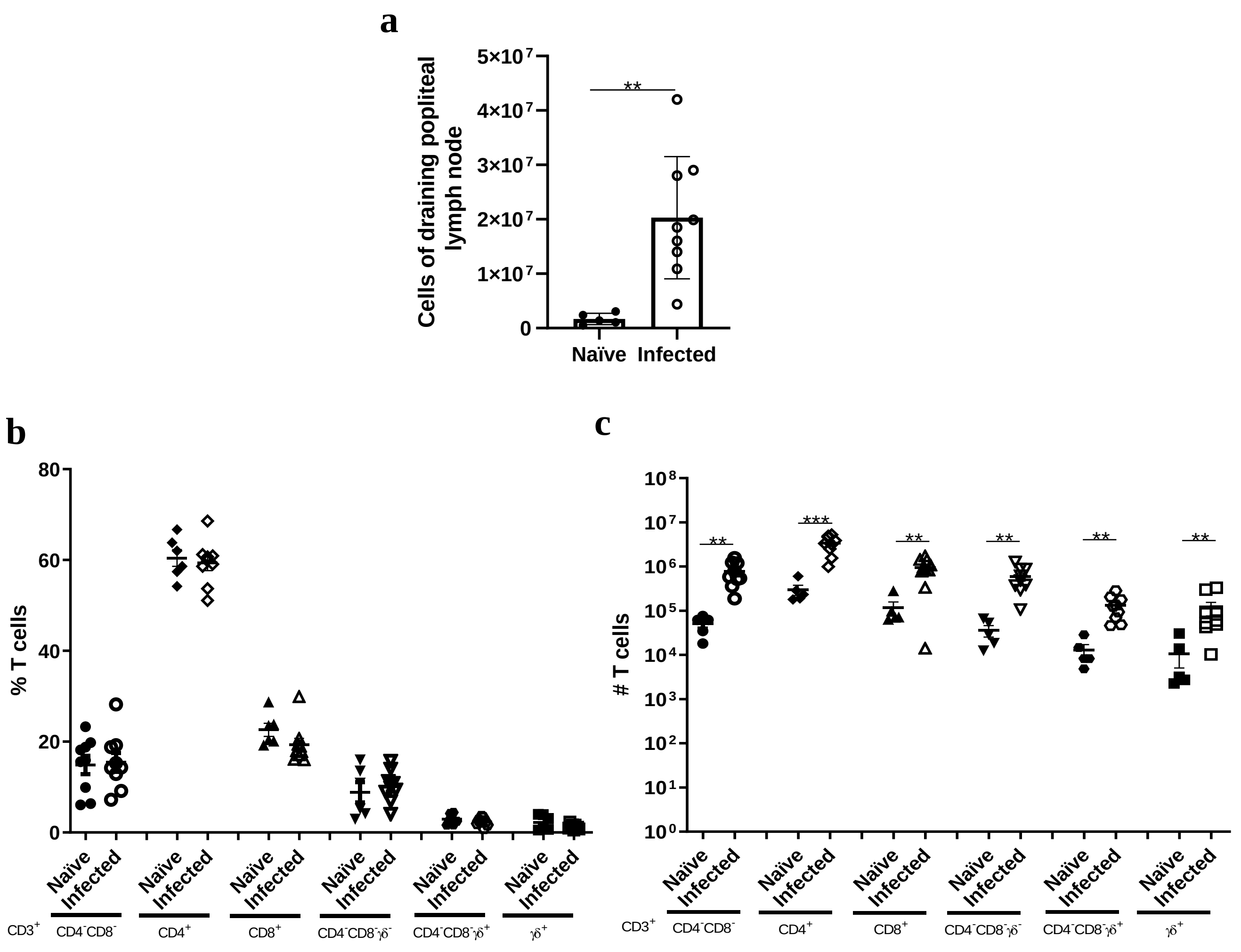
a
Cells of draining popliteal lymph node
5×10⁷
4×10⁷
3×10⁷
2×10⁷
1×10⁷
0
**
Naïve
Infected
b
% T cells
80
60
40
20
0
c
### T cells
10⁸
10⁷
10⁶
10⁵
10⁴
10³
10²
10¹
10⁰
**
***
Naïve
Infected
CD3⁺
CD4⁻CD8⁻
CD4⁺
CD8⁺
CD4⁻CD8⁻γδ⁻
CD4⁻CD8⁻γδ⁺
γδ⁺

Sup Fig 3

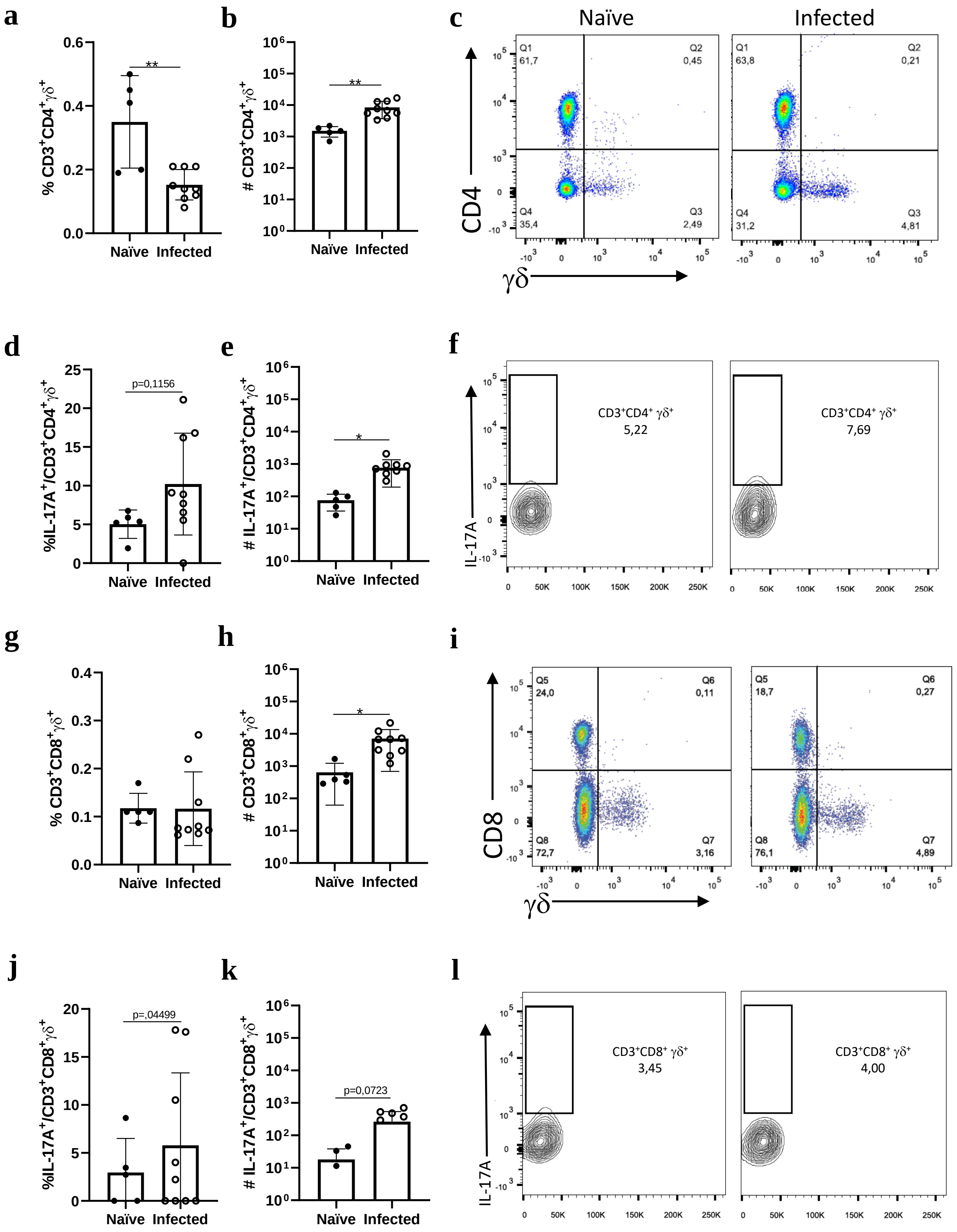
a
% CD3+CD4+γδ+
**
Naïve Infected
b
# CD3+CD4+γδ+
**
Naïve Infected
c
Naïve
Infected
CD4
γδ
Q1 61,7
Q2 0,45
Q4 35,4
Q3 2,49
Q1 63,8
Q2 0,21
Q4 31,2
Q3 4,81
d
%IL-17A+/CD3+CD4+γδ+
p=0,1156
Naïve Infected
e
# IL-17A+/CD3+CD4+γδ+
*
Naïve Infected
f
IL-17A
CD3+CD4+ γδ+ 5,22
CD3+CD4+ γδ+ 7,69
g
% CD3+CD8+γδ+
Naïve Infected
h
# CD3+CD8+γδ+
*
Naïve Infected
i
CD8
γδ
Q5 24,0
Q6 0,11
Q8 72,7
Q7 3,16
Q5 18,7
Q6 0,27
Q8 76,1
Q7 4,89
j
%IL-17A+/CD3+CD8+γδ+
p=,04499
Naïve Infected
k
# IL-17A+/CD3+CD8+γδ+
p=0,0723
Naïve Infected
l
IL-17A
CD3+CD8+ γδ+ 3,45
CD3+CD8+ γδ+ 4,00

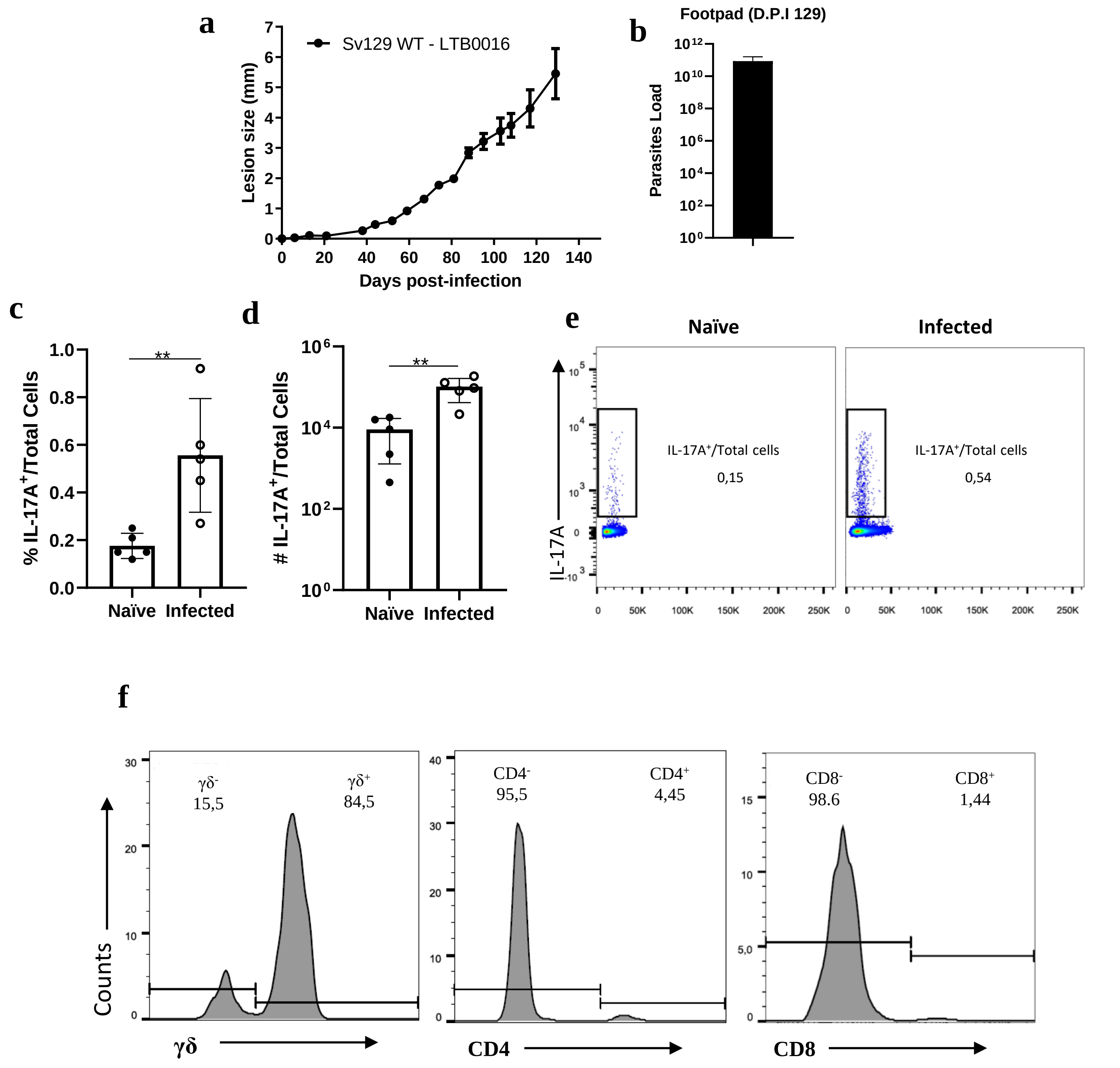
a
Sv129 WT - LTB0016
Lesion size (mm)
Days post-infection
b
Footpad (D.P.I 129)
Parasites Load
c
% IL-17A+/Total Cells
**
Naïve
Infected
d
### IL-17A+/Total Cells
**
Naïve
Infected
e
Naïve
Infected
IL-17A
IL-17A+/Total cells
0,15
IL-17A+/Total cells
0,54
f
Counts
γδ-
15,5
γδ+
84,5
γδ
CD4-
95,5
CD4+
4,45
CD4
CD8-
98.6
CD8+
1,44
CD8

Sup Fig 5

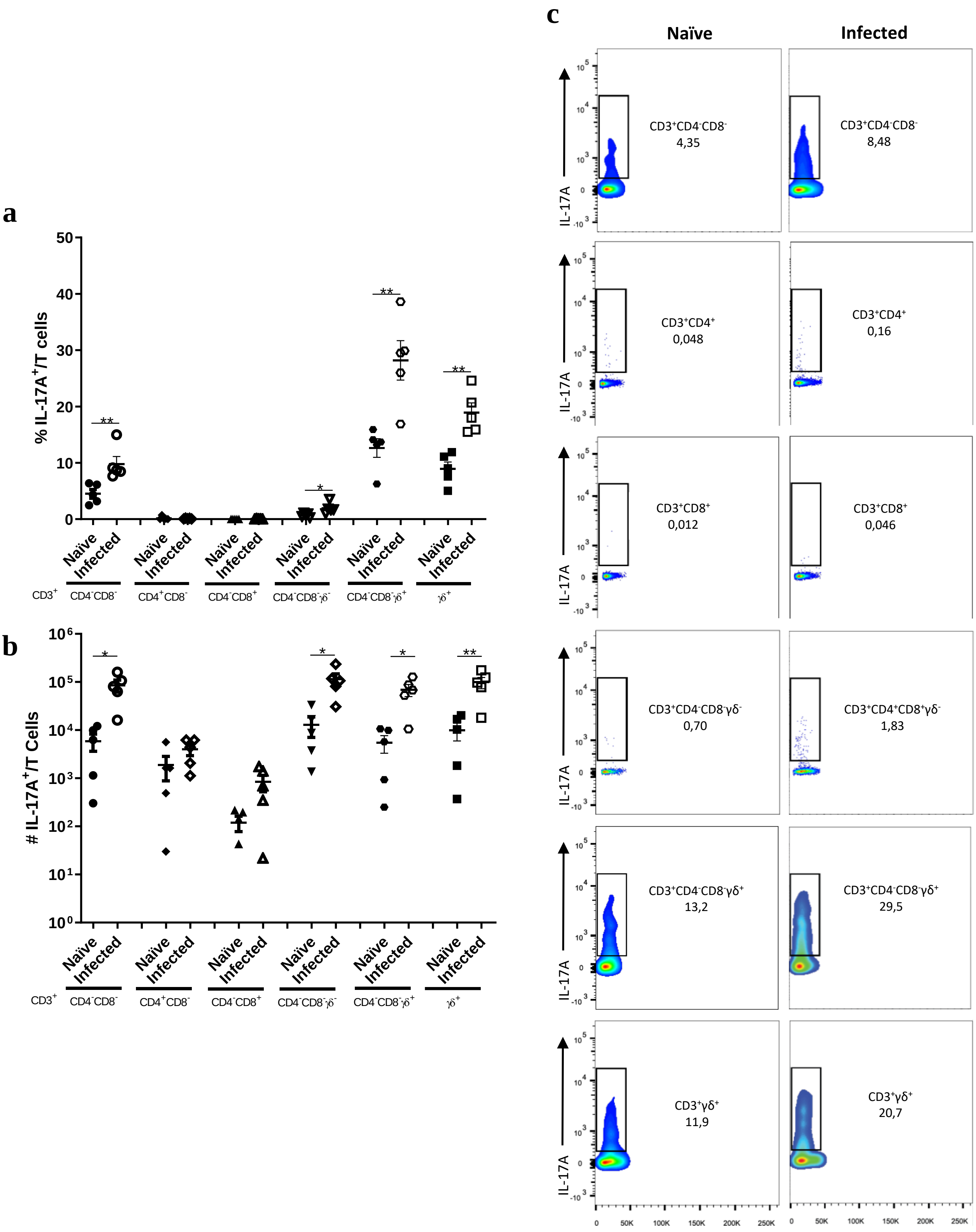
a
% IL-17A+/T cells
**
**
**
*
Naïve
Infected
CD3+
CD4-CD8-
CD4+CD8-
CD4-CD8+
CD4-CD8-γδ-
CD4-CD8-γδ+
γδ+
b
### IL-17A+/T Cells
c
Naïve
Infected
IL-17A
CD3+CD4-CD8-
4,35
CD3+CD4-CD8-
8,48
CD3+CD4+
0,048
CD3+CD4+
0,16
CD3+CD8+
0,012
CD3+CD8+
0,046
CD3+CD4-CD8-γδ-
0,70
CD3+CD4+CD8+γδ-
1,83
CD3+CD4-CD8-γδ+
13,2
CD3+CD4-CD8-γδ+
29,5
CD3+γδ+
11,9
CD3+γδ+
20,7

Sup Fig 6

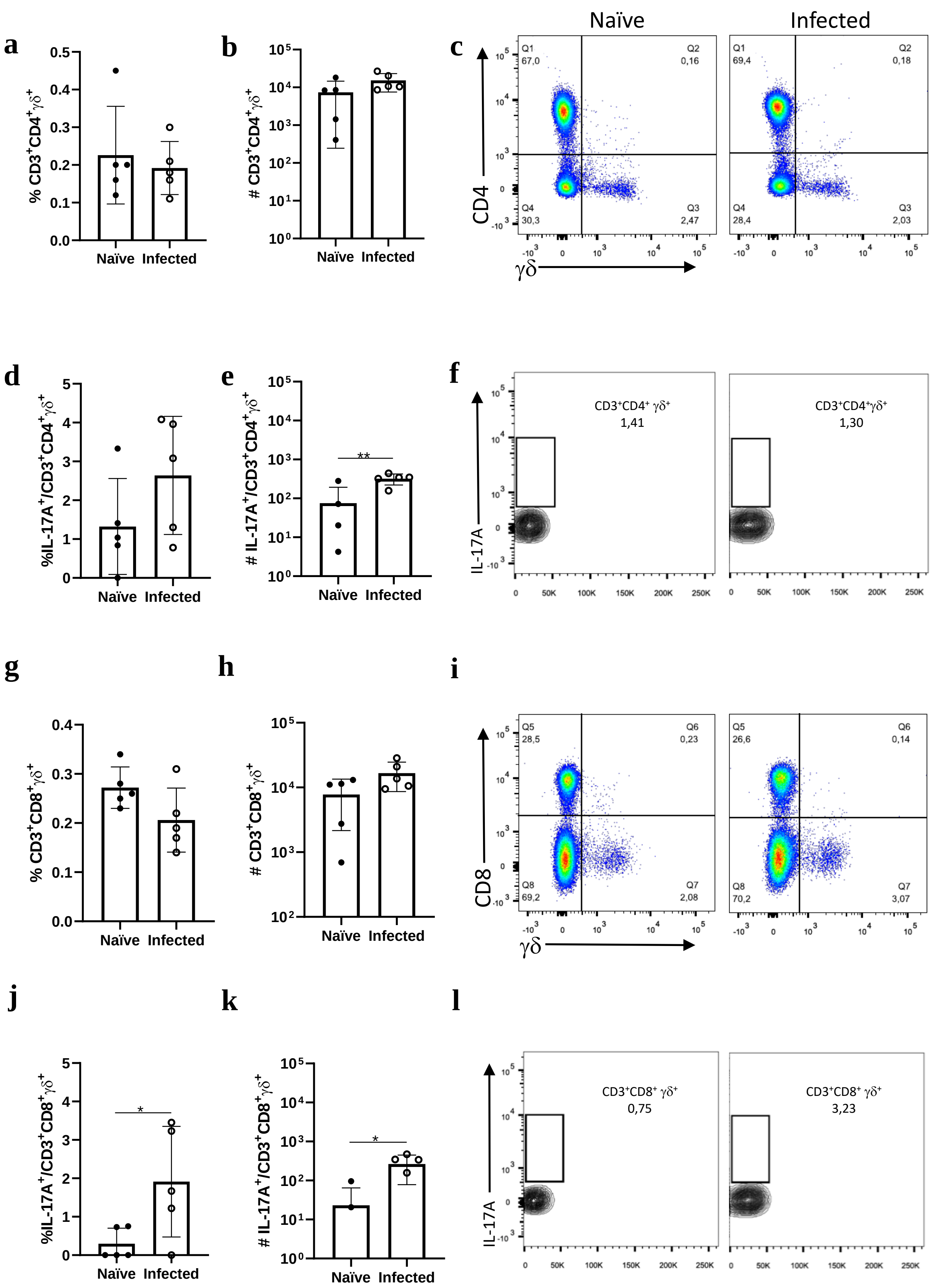

a
b
c
Naïve
Infected
d
e
f
g
h
i
j
k
l
CD3⁺CD4⁺ γδ⁺
1,41
1,30
CD3⁺CD8⁺ γδ⁺
0,75
3,23

Sup Fig 7

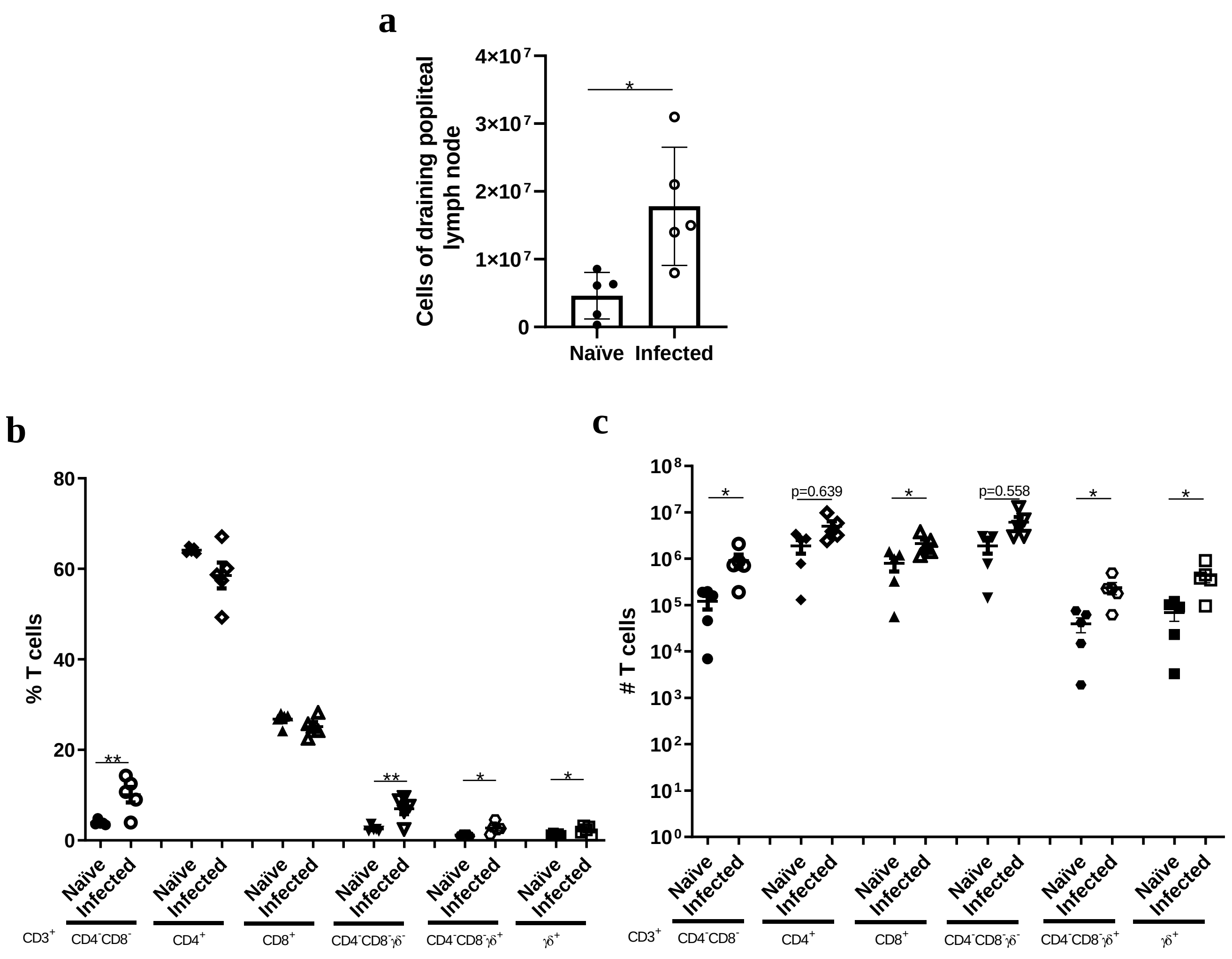
a
Cells of draining popliteal lymph node
4×10⁷
3×10⁷
2×10⁷
1×10⁷
0
*
Naïve
Infected
b
% T cells
80
60
40
20
0
**
**
*
*
Naïve
Infected
CD3+
CD4-CD8-
CD4+
CD8+
CD4-CD8-γδ-
CD4-CD8-γδ+
γδ+
c
### T cells
10⁸
10⁷
10⁶
10⁵
10⁴
10³
10²
10¹
10⁰
*
p=0.639
*
p=0.558
*
*
Naïve
Infected
CD3+
CD4-CD8-
CD4+
CD8+
CD4-CD8-γδ-
CD4-CD8-γδ+
γδ+

Sup Fig 8

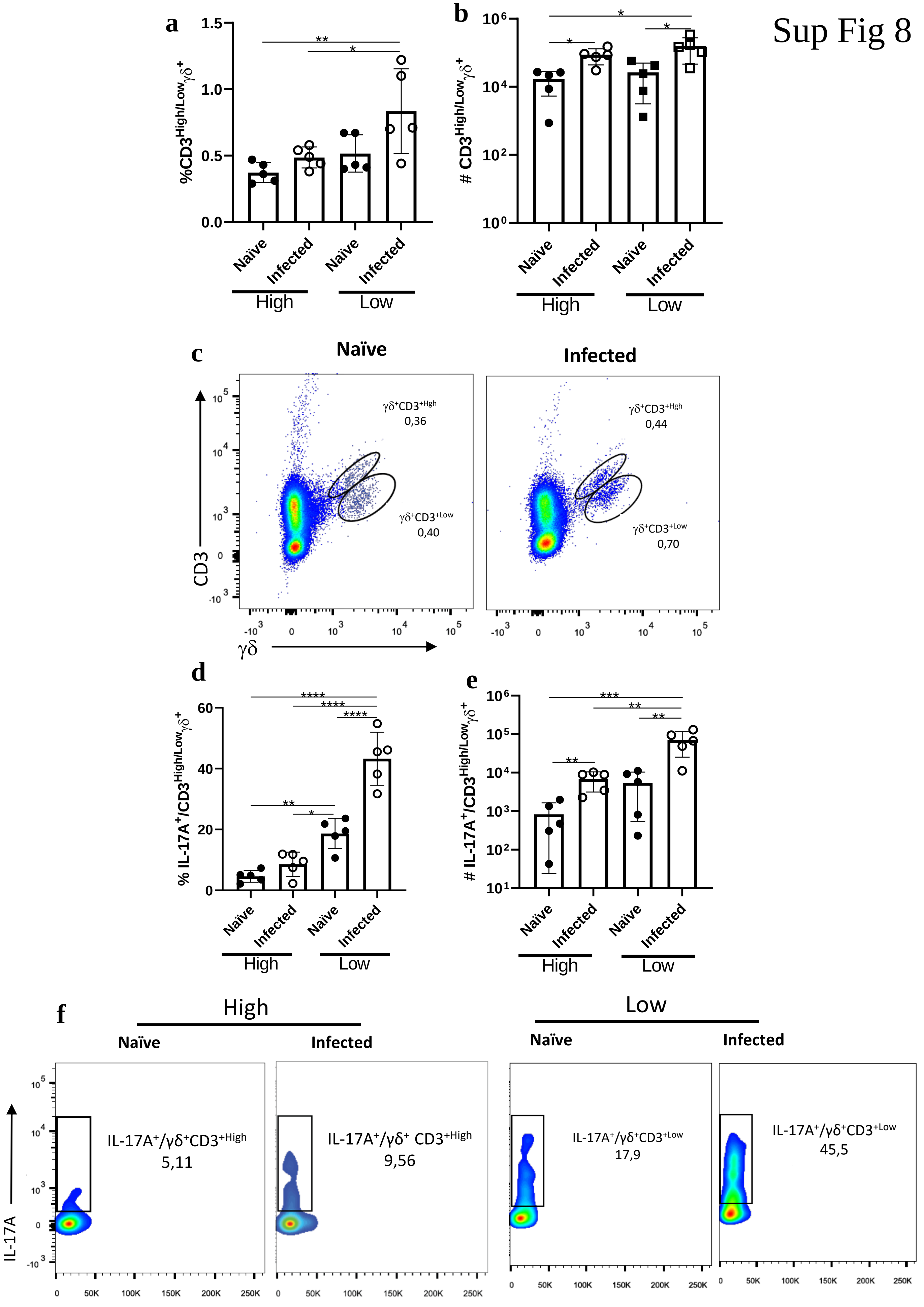
a
%CD3High/Low γδ+
1.5
1.0
0.5
0.0
**
*
Naïve
Infected
Naïve
Infected
High
Low
b
### CD3High/Low γδ+
10⁶
10⁴
10²
10⁰
c
Naïve
Infected
γδ+CD3+Hgh
0,36
γδ+CD3+Low
0,40
γδ+CD3+Hgh
0,44
γδ+CD3+Low
0,70
CD3
γδ
d
% IL-17A+/CD3High/Low γδ+
60
40
20
0
****
**
e
### IL-17A+/CD3High/Low γδ+
***
f
High
Low
Naïve
Infected
IL-17A+/γδ+CD3+High
5,11
IL-17A+/γδ+ CD3+High
9,56
IL-17A+/γδ+CD3+Low
17,9
IL-17A+/γδ+CD3+Low
45,5
IL-17A
0
50K
100K
150K
200K
250K

Sup Fig 9

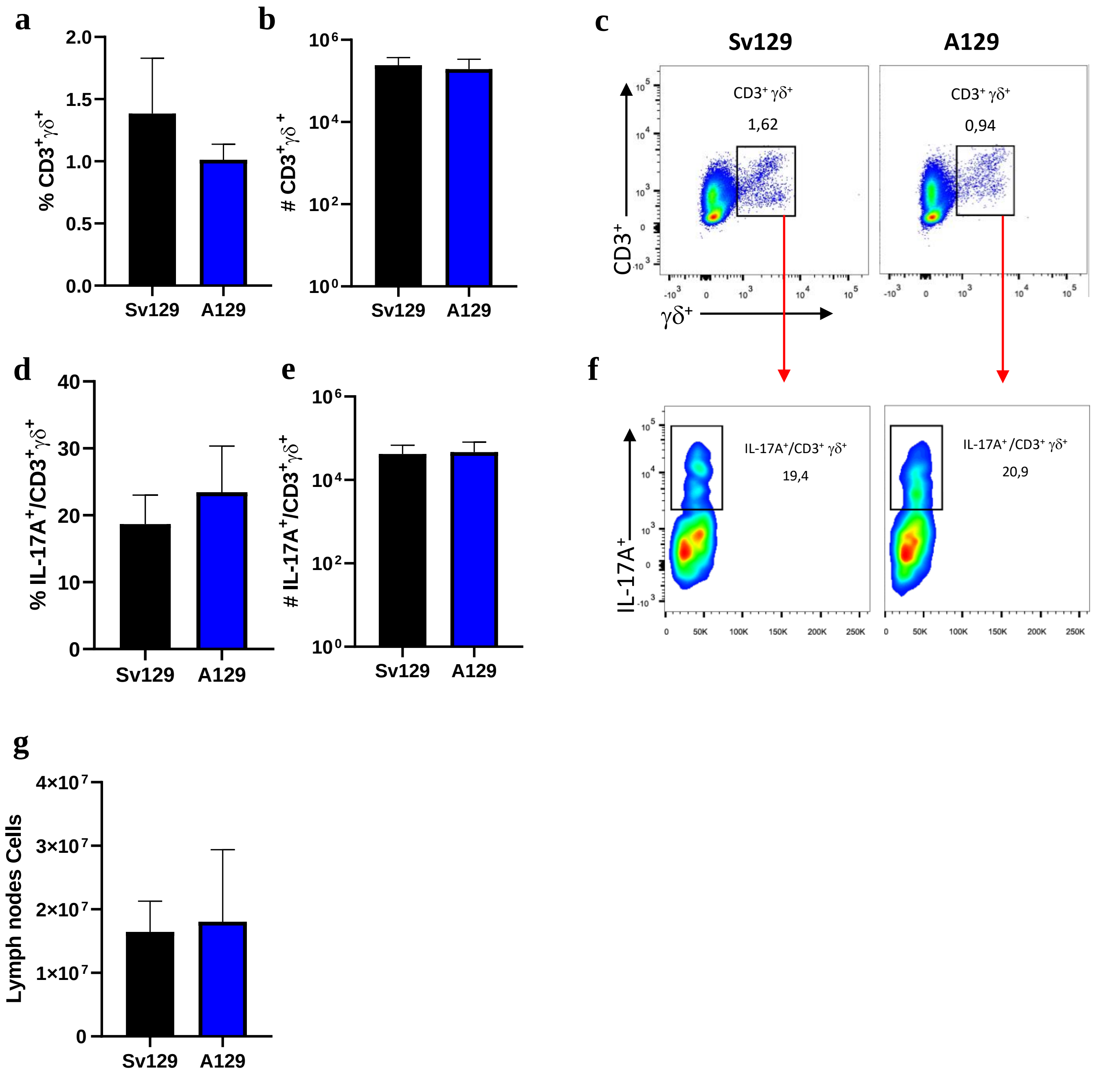

a
2.0
1.5
1.0
0.5
0.0
% CD3+γδ+
Sv129
A129
b
10⁶
10⁴
10²
10⁰
# CD3+γδ+
Sv129
A129
c
Sv129
A129
CD3+ γδ+
1,62
CD3+ γδ+
0,94
CD3+
γδ+
d
40
30
20
10
0
% IL-17A+/CD3+γδ+
Sv129
A129
e
# IL-17A+/CD3+γδ+
Sv129
A129
f
IL-17A+/CD3+ γδ+
19,4
IL-17A+/CD3+ γδ+
20,9
IL-17A+
0
50K
100K
150K
200K
250K
g
4×10⁷
3×10⁷
2×10⁷
1×10⁷
0
Lymph nodes Cells
Sv129
A129

Sup Fig 10

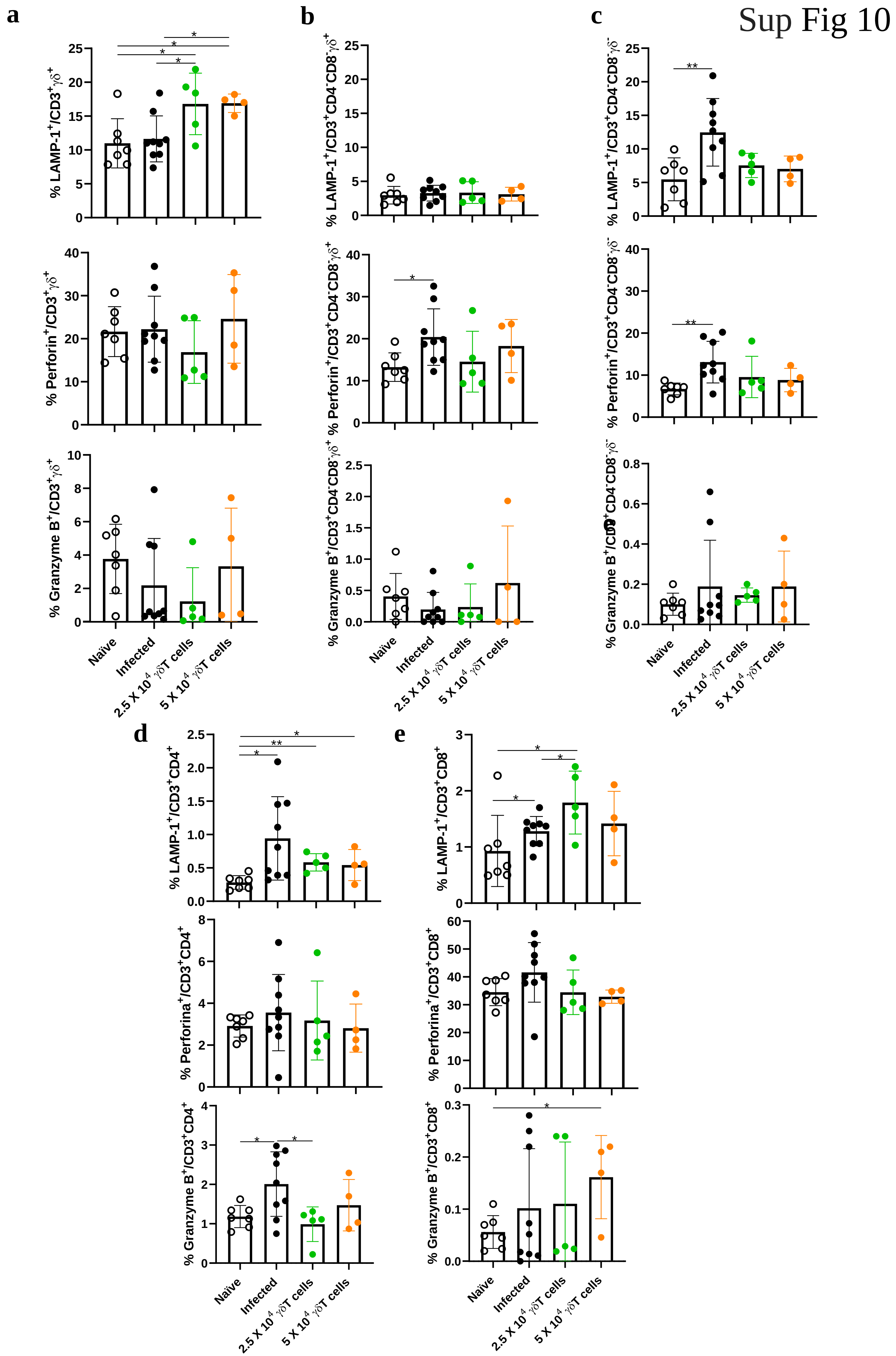

a
b
c
d
e
% LAMP-1+/CD3+γδ+
% Perforin+/CD3+γδ+
% Granzyme B+/CD3+γδ+
% LAMP-1+/CD3+CD4-CD8-γδ+
% Perforin+/CD3+CD4-CD8-γδ+
% Granzyme B+/CD3+CD4-CD8-γδ+
% LAMP-1+/CD3+CD4-CD8-γδ-
% Perforin+/CD3+CD4-CD8-γδ-
% Granzyme B+/CD3+CD4-CD8-γδ-
% LAMP-1+/CD3+CD4+
% Perforina+/CD3+CD4+
% Granzyme B+/CD3+CD4+
% LAMP-1+/CD3+CD8+
% Perforina+/CD3+CD8+
% Granzyme B+/CD3+CD8+
Naïve
Infected
2.5 X 10^4 γδT cells
5 X 10^4 γδT cells

a

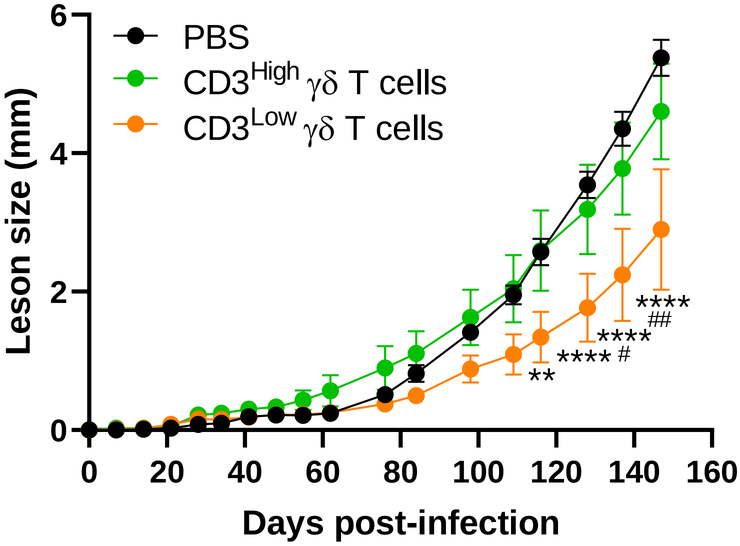
PBS
CD3High γδ T cells
CD3Low γδ T cells
Leson size (mm)
Days post-infection
0
2
4
6
0
20
40
60
80
100
120
140
160
**
****
#
****
****
##

b

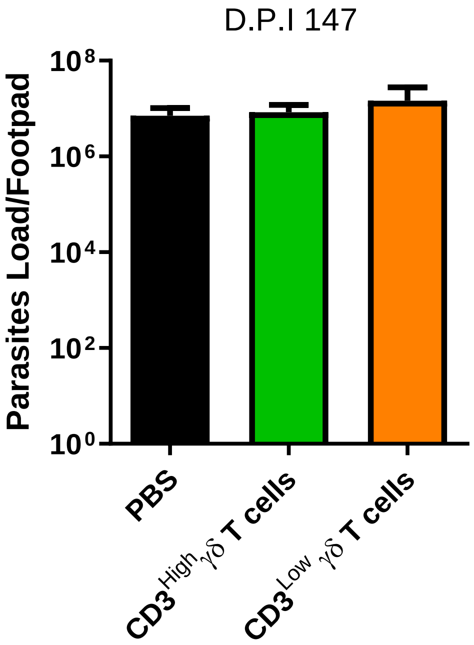
D.P.I 147
Parasites Load/Footpad
10⁸
10⁶
10⁴
10²
10⁰
PBS
CD3High γδ T cells
CD3Low γδ T cells
